# BRCA1-BARD1 regulates transcription through modulating topoisomerase IIβ

**DOI:** 10.1101/2020.12.12.422337

**Authors:** Jaehyeon Jeong, Keunsoo Kang, Doo Sin Jo, Anh TQ Cong, Donguk Kim, Dong-Hyung Cho, Matthew J Schellenberg, Stuart K. Calderwood, Benjamin P.C. Chen, Heeyoun Bunch

## Abstract

RNA polymerase II (Pol II)-dependent transcription in stimulus-inducible genes requires topoisomerase IIβ (TOP2B)-mediated DNA strand break and the activation of DNA damage response signaling in humans. Here, we report a novel function of the <u>br</u>east <u>ca</u>ncer 1 (BRCA1)-<u>B</u>RCA1 <u>a</u>ssociated <u>r</u>ing <u>d</u>omain <u>1</u> (BARD1) complex, in this process. We found that BRCA1 is phosphorylated at S1524 by the kinases ATM and ATR during gene activation and that this event is essential for productive transcription. Our *in vitro* biochemical analyses showed TOP2B and BARD1 interaction and colocalization in the *EGR1* transcription start site (TSS) and that the BRCA1-BARD1 complex ubiquitinates TOP2B, which appears to stabilize TOP2B protein in the cell and binding to DNA. Intriguingly, BRCA1 phosphorylation at S1524 controls this interaction. In addition, genomic analyses indicated colocalization between TOP2B and BRCA1 in a large number of protein-coding genes. Together, these findings reveal the novel function of the BRCA1-BARD1 complex in gene expression and in the regulation of TOP2B during Pol II transcription.

**Significance Statement:** Maintaining genomic integrity against cellular and extracellular genotoxic elements is essential for normal cell growth and function. Recent studies indicated that stimulus-induced transcription provokes topoisomerase IIβ-mediated DNA strand break and DNA damage response signaling, requiring DNA repair to be coupled with transcription. Here, we present a novel role for the BRCA1-BARD1 complex in regulating the transcription of serum-inducible genes and the stability of topoisomerase IIβ. The mechanism involving topoisomerase IIβ ubiquitination by the BRCA1-BARD1 complex and the phosphorylation of BRCA1 S1524 upon transcriptional activation appears to function as a switch to the reaction. Our findings provide the first evidence of functional interaction between the BRCA1-BARD1 complex and topoisomerase IIβ in transcription in humans.

## Introduction

Transcription is the first and most immediate step in gene expression, but it also contributes to genomic instability. The innate genotoxic mechanisms appearing during transcription include R-loop-mediated susceptibility of the non-template DNA and the collision of replication-transcription machineries on chromosomes, which result in single-strand or <u>d</u>ouble-<u>s</u>trand <u>b</u>reakage (DSB) of DNA (Costantino and Koshland, 2018; Gomez-Gonzalez and Aguilera, 2019). In addition, a few studies in the past decade have reported the formation of DSB, a so-called programed DNA break, during transcription, in which DSB is induced by stimuli such as steroid-hormone receptor binding and neurotransmitter- and serum-induced gene activation (Bunch et al., 2015; Bunch et al., 2014; Haffner et al., 2010; Ju et al., 2006; Madabhushi et al., 2015; Puc et al., 2015). In the latter cases, TOP2B is responsible for generating the DSB (Bunch et al., 2015; Ju et al., 2006; Madabhushi et al., 2015). It has been suggested that <u>TO</u>P2B-mediated <u>DSB</u> (Top-DSB) is persistent rather than instantaneously repaired, which is expected in typical topoisomerase-mediated topological resolution (Pommier et al., 2016). This speculation is based on the fact that the catalytic activity of TOP2B during transcriptional activation induces <u>D</u>NA <u>d</u>amage <u>r</u>esponse (DDR) signaling (Bunch, 2017; Pommier et al., 2016). Importantly, DSB is required for efficient transcription because inhibiting either TOP2B or major DDR kinases, including <u>DNA</u>-dependent <u>p</u>rotein <u>k</u>inase (DNA-PK) and <u>a</u>taxia telangiectasia <u>m</u>utated (ATM), has been shown to deregulate Pol II translocation from initiation to processive elongation; this in turn deregulates productive RNA synthesis in *HSP70*, enhancer RNA, and the serum- and estrogen receptor-activated genes in humans (Bunch et al., 2015; Bunch et al., 2014; Ju et al., 2006; Madabhushi et al., 2015; Puc et al., 2015). These observations suggest a potential innate cause for genomic aging or mutability attributed to gene expression during transcription, an inevitable process in growing, differentiating, and maintaining the cells and organs of any organism. Therefore, the DNA repair system, coupled with transcription, plays a crucial and essential role in protecting the genome from accumulating instability.

Repair of most genomic DSBs involves two major pathways, the <u>n</u>on-<u>h</u>omologous <u>e</u>nd joining (NHEJ) and <u>h</u>omology-<u>d</u>irected repair (HDR)(Chapman et al., 2012). NHEJ and HDR are executed by different protein repair factors, namely 53BP1 and BRCA1, respectively (Anantha et al., 2017; Jayavaradhan et al., 2019). Because of the requirement for an intact homologous strand, HDR is thought to be more restricted in higher eukaryotes and is prevalent in the S/G_2_ cell cycle phase, while NHEJ predominates in the G_1_ phase of the cell cycle (Bau et al., 2006; Mao et al., 2008). However, BRCA1 appears to be involved in NHEJ as well and reportedly participates in precise NHEJ during the G_1_ phase (Jiang et al., 2013). In addition, recent studies have found that BRCA1 is recruited to R-loop and Pol II pausing sites (Chiang et al., 2019; Hatchi et al., 2015; Herold et al., 2019) and plays critical roles in removing and repairing TOP2-DNA adducts in replication and estrogen-induced transcription (Aparicio et al., 2016; Morimoto et al., 2019; Sasanuma et al., 2018). The expression of BRCA1 is maintained at a relatively low level in the cell, and BRCA1 activity is regulated by post-translational modifications, including phosphorylation, methylation, sumoylation, and ubiquitination (https://www.uniprot.org/uniprot/P38398). Upon DNA damage, BRCA1 is phosphorylated at multiple residues, including serines 1387, 1423, 1457, and 1524, by ATM (Cortez et al., 1999; Gatei et al., 2000). BRCA1 often functions in complex with BARD1 as an E3 ubiquitin ligase during DNA repair and cell proliferation (Baer and Ludwig, 2002; Densham and Morris, 2017; Starita et al., 2005; Stewart et al., 2018; Verma et al., 2019). BRCA1-BARD1 heterodimers are auto-ubiquitinated, modifications which provokes the ubiquitin ligase activity of this protein complex (Mallery et al., 2002). So far, H2A, NF2, and estrogen receptor *α* are known ubiquitination substrates for the BRCA1-BARD1 complex (Ma et al., 2010; Stewart et al., 2018; Verma et al., 2019).

Transcription shows marked activity in G_1_ phase to mediate the expression of the proteins and biomolecules required for genome replication in the S phase and later steps in cell proliferation. This process is initiated by <u>h</u>uman <u>i</u>mmediate <u>e</u>arly <u>g</u>enes (hIEGs), which are transcribed as cells transit from the G_0_ to the G_1_ phase. Many such genes are transcription factors and proto-oncogenes, such as *JUN, FOS, MYC*, and *EGR1*. These genes are expressed rapidly, upon the receipt of cell proliferation signals, and their transcription requires the mechanism underlying Pol II promoter-proximal pausing (Pol II pausing), followed by pause release (Bunch et al., 2015; Liu et al., 2015). In the resting state of transcription, Pol II pausing occurs at approximately +25 to +100 from the <u>t</u>ranscription <u>s</u>tart <u>s</u>ite (TSS) of a large number of protein-coding and non-coding genes in metazoan cells (Adelman and Lis, 2012; Benjamin and Gilmour, 1998; Bunch et al., 2016; Bunch et al., 2014; Core et al., 2008; Rahl et al., 2010). Gene activation involves the release of Pol II from the pausing site and the resumed production of a full-length transcript (Bunch et al., 2014; Chen et al., 2015; Peterlin and Price, 2006; Rahl et al., 2010; Zobeck et al., 2010). It has been reported that Top-DSB and DDR signaling are accompanied by and required for Pol II pause release and gene activation in hIEGs (Bunch, 2016; Bunch et al., 2015; Madabhushi et al., 2015). This finding raises important questions about how TOP2B and Top-DSB are regulated and repaired during transcription.

In this study, we investigated whether BRCA1 regulates the transcription of stimulus-inducible genes and, in particular, its function in the regulation of TOP2B during transcriptional pausing and activation. Through biochemical and cell-based analyses, we found that the BRCA1-BARD1 complex is essential for the expression of the *EGR1* gene, a representative hIEG that utilizes Pol II pausing for gene regulation. Upon serum induction, BRCA1 is phosphorylated at S1524 by ATM and ATR in hIEGs, an important requirement for active transcription. Our data indicate that TOP2B binds to a fragment including –132 to +62 with a strongest affinity (K_d_ = 59.9 ± 6.1 nM) in *EGR1* TSS and BARD1 interacts with TOP2B within the DNA segment of 119 bp (–132 to –15). Intriguingly, ubiquitination of TOP2B enhances binding to these segments while the deubiquitinated form cannot. In addition, our biochemical analyses suggest that the phosphorylation status of BRCA1 at S1524 appears to control the BARD1-TOP2B interaction: the BRCA1 S1524A mutant facilitates TOP2B ubiquitination by BRCA1-BARD1 complex, which leads to a stronger association between TOP2B and *EGR1* TSS. Consistently, WT or a phospho-mimetic BRCA1 mutant, S1524D, which weakens TOP2B ubiquitination, destabilizes TOP2B binding to the *EGR1* TSS. We also find that the BRCA1-BARD1 complex is important for TOP2B stabilization as BARD1 KD reduces TOP2B protein levels. Consistently, genomic analyses showed the colocalization of BRCA1 and TOP2B in a large number of protein-coding genes in humans, suggesting their widespread functional involvement. Together, these results suggest a novel role for BRCA1-BARD1 complex in stress-inducible gene transcription by regulating TOP2B stability during transcription activation.

## Materials & Methods

### Cell culture and experimental conditions

HEK293 cells were grown in complete medium, composed of DMEM (Corning), supplemented with 10% FBS (Gibco) and 1% <u>p</u>enicillin/<u>s</u>treptomycin solution (P/S, Gibco). For serum-induction experiments, HEK293 cells were grown to about 80% confluence. The cells were incubated in DMEM, including 0.1% FBS and 1% P/S solution, for 17.5 h and then were induced using serum through incubation in DMEM, supplemented with 18% FBS and 1% P/S solution. After serum induction, cells were collected at the corresponding time points listed in the figures. For the ATM and ATR inhibitor experiment, HEK293 cells were incubated in the 0.1% serum media for 17.5 h. The media were exchanged with the 0.1% serum media including KU55933 (Abcam, ab120637), VE821 (Sigma, SML-1415), or caffeine (Sigma, C0750) at a final concentration of 10 µM, 1 μM, or 3 mM in 0.1% DMSO (for KU55933 and VE821) and 4% water (for caffeine) of the total media volume. The cells were incubated for 1 h before serum induction for 15 min with 18% serum media, including the chemicals at the same final concentration. Control cells were prepared side-by-side using DMSO only at the same final concentration. Stock solutions were made by dissolving kinase inhibitors as 10 mM, 1 mM, and 75 mM in DMSO (KU55933 and VE821) or water (caffeine) to target the final concentrations of 10 μM, 1 μM, and 3 mM for KU55933, VE821, and caffeine.

### Cell transfection

HEK293 cells were grown to approximately 70% confluence in complete media. The media were exchanged with the complete media without antibiotics immediately before transfecting the cells with scrambled (#6568, Cell Signaling) or BARD1 siRNA duplexes (SR300400, Origene) and BRCA1 targeting siRNA species (sc-29219, Santa Cruz Biotechnology) dissolved in serum-free DMEM using Lipofectamine 2000 (Invitrogen) according to the manufacturer’s instructions. The cells were collected after 48 h or 72 h incubation for RNA or protein analyses, respectively, or were subjected to serum starvation/induction for chromatin immunoprecipitation assays, Western blotting, or immunoprecipitation.

### RNA quantification

RNA molecules were extracted using RNeasy kit (Qiagen) following the manufacturer’s instructions. From each experimental condition, 0.6 or 1 μg extracted RNA was converted into cDNA by reverse transcription using a High Capacity cDNA Reverse Transcription kit (Promega A3500) or a ReverTra Ace qPCR RT Master Mix (Toyobo). RT-qPCR was conducted with equal amounts of resultant cDNAs and indicated primers (Table S1) using Platinum Tag DNA Polymerase High Fidelity (Invitrogen) under thermal cycling for 2 min at 94°C followed by 25 cycles of 20 s at 94°C, 30 s at 55°C, and 30 s at 68°C or through GO taq polymerase (Promega) for 2 min at 95°C followed by 25 cycles of 30 s at 95°C, 30 s at 55°C, and 30 s at 72°C. Real-time quantitative PCR (qRT-PCR) was performed with a CFX96 Real-Time PCR Detection System (Bio-Rad) or Thermal Cycler Dice Real Time System III (Takara) using β-actin as an internal control. SYBR Green Realtime PCR master mix was purchased from Toyobo. The primers used for the study are listed in Table S1. The results are presented as means SEMs after normalization by the sham (no serum, DMSO) group.

### DNA templates

The DNA template of *EGR1* promoter and early transcript, including –432 to +323, was amplified from HeLa <u>n</u>uclear <u>e</u>xtract (NE) using a pair of primers (Table S1). The amplified product was cloned into a pCR-Blunt-TOPO plasmid, called pTOPO-EGR1. The biotinylated template was generated via PCR using the cloned vector as a template and a set of primers, one conjugated with biotin at the 5′ end (Table S1). The PCR product was sequence-verified, gel-extracted, and purified using the Qiaquick gel extraction kit (Qiagen) before further experimentation. The mutant pTOPO-EGR1 vectors, including the altered sequences between +141 and +160 and a point-substitution at +69 that generated a SfoI site were synthesized via Quikchange site-directed mutagenesis, using pairs of primers (Table S1) incorporating these mutations. All primers that were designed and used in the study were purchased from Integrated DNA Technology, and their sequences are listed in Table S1.

### Proteins, expression vectors, and purification

A full-length BRCA1 was amplified from a BRCA1 expression vector, pcBRCA1-385, provided by Dr. Mike Erdos at the National Institutes of Health and cloned into bacterial expression vector pET17b, including an His^6^ tag, with a pair of primers (Table S1) to produce pET-WT-BRCA1. The pET17b vectors expressing mutant BRCA1 species S1524A and S1524D (pET-SA-BRCA1 and pET-SD-BRCA1) were generated by Quikchange site-directed mutagenesis, using pairs of primers (Table S1) that incorporated the desired amino-acid substitutions. A full-length coding sequence of BARD1 was amplified from BARD1 cDNA (HG15850-CH, Sino Biological) and cloned into another bacterial expression vector, pET29b including His^6^ tag in the C-terminal. A full length coding sequence of TOP2B was amplified from our cDNA library of HEK293 cells. Our bacterial-derived TOP2B construct encoded the N-terminal 566 amino acids. It was cloned to pET21a, including His^6^ in the N-terminal domain. The primers used to amplify and clone the coding sequences of BARD1 and TOP2B are listed in Table S1. These five protein expression vectors were sequence-verified, and each was transformed into BL21. The His^6^-taged proteins were purified using Ni beads (Invitrogen). Protein expression was induced at 20°C for 16 h using 0.5–1 mM IPTG as a final concentration (Figure S1). The *E. coli* cells expressing WT-, SA-, and SD-BRCA1 were harvested and lysed with the xTractor bacterial cell lysis buffer (Clontech) and sonication. The Ni wash buffers included 10 mM and 25 mM imidazole, 0.5 M NaCl, 10 mM Tris-HCl pH 7.6, 0.5% Triton X-100, and 10% glycerol. The wash was performed-twice with 10 mM imidazole wash buffer, and then twice with 25 mM buffer for BRCA1 and four times with 25 mM buffer for BARD1 and TOP2B. Bead-bound proteins were eluted with 500 mM imidazole buffer including 0.02% NP40. The eluted proteins were subjected to SDS-PAGE and immunoblotting to be verified. All protein purification buffers included freshly added protease inhibitors, 1 mM benzamidine (Sigma), 0.25 mM PMSF (Sigma), aprotinin (Sigma A6279, 1:1000), and 1 mM Na-metabisulfite (Sigma). DNA encoding full-length human TOP2B (amino acids 1-1621) was amplified by PCR from pcDNA6.2/C-YFPDest-TOP2β (Schellenberg et al., 2017) with primers that incorporate a HRV3C protease site at the N-terminus of TOP2β and cloned into the vector pmCentr2 using ligation-independent cloning. LR Clonase II (Thermofisher) was used to transfer the HRV3C-TOP2B insert into pcDNA6.2/N-YFPDest to generate the YFP-TOP2B plasmid, transformed into the Stbl3 *E. coli* strain (Thermofisher), and purified using a Plasmid DNA Gigaprep kit (Zymo). The insert was fully sequenced to confirm the absence of mutations (see Table S1 for primer sequences). YFP-TOP2B was transfected into HEK293F cells in Hyclone TransFx media (Cytiva) using PEI (Polysciences) and purified using the YFP-tag system (Schellenberg et al., 2018). HEK293F cell pellets were lysed in 36 mL lysis buffer [50 mM Tris-HCl pH 8.0, 400 mM NaCl, 0.1% (v/v) NP-40 substitute (Sigma), and 1 mM TCEP] supplemented with Complete-EDTA free protease inhibitor cocktail (Roche), and sonicated with 3 cycles of 5 s sonication using a Branson sonicator set to 55% power, followed by 30 s cooling periods. Crude lysate was centrifuged at 25,000x g for 10 minutes, then passed over a column with pre-equilibrated anti-GFP/YFP sepharose resin. The color of the resin was monitored visually, and additional resin was used if the resin became saturated. The resin was washed 6 times with lysis buffer (supplemented with 100 nM USP2 (purified in-house from Addgene plasmid 36894) for de-ubiquitinated TOP2B), then 3 times in ATP wash buffer (50mM Tris-HCl pH 8.0, 200mM NaCl, 2mM MgCl_2_, 0.05% Tween-20, 1mM TCEP, add 2mM ATP; with 100 nM USP2 for de-ubiquitinated TOP2B) followed by a 15 minute incubation at room temperature to remove HSP70 contaminant. Resin was then washed 3 times with cold size-exclusion buffer (20 mM Tris-HCl pH 7.5, 500 mM NaCl, 1 mM TCEP), then 1.5 column volumes of size-exclusion buffer supplemented with 45 μg/mL HRV3C protease. Column was capped tightly and incubated overnight to cleave TOP2B from the YFP-tag. TOP2B was eluted the next day by washing the resin with size-exclusion and diluted in 3 volumes of low salt buffer (20 mM Tris-HCl pH 7.5, 1 mM DTT), loaded onto a 6mL SOURCE 15S column (GE Healthcare), and eluted with a linear gradient of 0-50% high salt buffer (20 mM Tris-HCl pH 7.5, 1 M NaCl). Fractions containing TOP2B were pooled and concentrated to 2 mL volume by ultrafiltration (Amicon) and run on a Superdex200 16/60 column (GE) in size-exclusion buffer. Fractions containing TOP2B were pooled, concentrated by ultrafiltration, and buffer exchanged into TOP2 storage buffer (20 mM HEPES pH 7.5, 500 mM NaCl, 1 mM TCEP, and 25% (v/v) glycerol. Protein aliquots were stored long term (>6 months) at –80°C or short term (<1 month) at –20°C.

### HeLa nuclear extract preparation

HeLa nuclei were provided by Taatjes laboratory (University of Colorado, Boulder). Using these nuclei, HeLa NE was generated. The nuclei were dissolved with 0.9 volumes of Buffer C (20 mM HEPES pH 7.9, 25% glycerol, 420 mM NaCl, 1.5 mM MgCl_2_, 0.2 mM EDTA) while stirring at 4°C. The mixture was dounced 20 times with pestle B before being stirred gently for 30 min at 4°C. Then the homogenized solution was centrifuged for 30 min at 13,000 rpm at 4°C. The supernatant was to dialyzed against Buffer D (20 mM HEPES pH 7.9, 20% glycerol, 100 mM KCl, 2 mM MgCl_2_, 0.2 mM EDTA) to conductivity 100–150 mM, which was measured by adding 25 µL sample to 5 mL HPLC water (Bunch et al., 2014). Buffers C and D included the fresh protease inhibitors described above and 1 mM DTT (complete protease inhibitors). The resulting HeLa NE was validated for protein quality and concentration by Western blotting for probing nuclear proteins, including RNA polymerase II and BRCA1, and Bradford assays to compare with a reference HeLa NE, provided by Taatjes laboratory.

### Immobilized template assay and transcription assay

The experimental procedure for immobilized template and transcription assays is identical to that of our previous report, except for the method of quantifying nascent RNAs synthesized *in vitro* (Bunch et al., 2014). Dynabeads M-280 Streptavidin (Invitrogen) was prepared with 2X B&W buffer (10 mM Tris-HCl, pH 7.5, 1 mM EDTA, 2 M NaCl) and incubated with biotin-conjugated *EGR1* template DNA (–432 to +323) at 10 ng DNA/μL beads. The template-conjugated beads were washed with 1X B&W buffer and 0.1 M Buffer D1 (20 mM HEPES, 20% glycerol, pH 7.6, pH 7.9, 0.1 mM EDTA, 100 mM KCl). Then 120 ng immobilized template was mixed with TF buffer (12.5 ng/μl dI-dC, 0.075% NP40, 5 mM MgCl_2_, 250 ng/µL BSA, 12.5 % glycerol, 100 mM KCl, 12.5 mM HEPES, pH 7.6, 62.5 μM EDTA, 10 µM ZnCl_2_) for pre-incubation with purified WT or mutant BRCA1 protein. The resultant template-protein complex was pulled-down using a magnet stand (Invitrogen) and resuspended in NE buffer (17.5 ng/µL dI-dC, 0.1% NP40, 7.5 mM MgCl_2_, 1.25 µg/μL BSA, 8.7% glycerol, 8.7 mM HEPES, pH 7.6, 44 μM EDTA, 130 mM KCl, 10 µM ZnCl_2_). HeLa NE and purified recombinant WT or mutant BRCA1 protein when indicated as T1 was added at 100 μg/reaction and incubated with agitation for 30 m at room temperature (RT) to assemble PIC. The template-protein complex was washed briefly with a 10 beads volume of TW buffer (13 mM HEPES, pH 7.6, 13% glycerol, 60 mM KCl, 7 mM MgCl_2_, 7 mM DTT, 100 μM EDTA, 0.0125% NP40, 10 μM ZnCl_2_) and then resuspended in Transcription Buffer I (13 mM HEPES, pH 7.6, 13% Glycerol, 60 mM KCl, 7 mM MgCl_2_, 10 μM ZnCl_2_, 7 mM DTT, 100 μM EDTA, 15 ng/μL dI-dC, 10 mM creatine phosphate). For transcription assay, a mixture of NTP in final concentrations of 250 µM A/G/C/U was added to initiate Pol II to polymerize mRNA molecules at 30°C. When T2 was indicated, purified WT or mutant BRCA1 protein was introduced, both during the PIC assembly and after 3 min of NTP addition. The polymerization reaction was allowed for 30 min total incubation time. Then, 1.5 Kunitz units DNase I (Qiagen) was added to the reaction and allowed to sit for an additional 15 min to remove the template DNA. All reactions were terminated with 5 volumes 1.2 X Stop buffer (0.6 M Tris-HCl, pH 8.0, 12 mM EDTA, 100 µg/mL tRNA). The pellet fraction, including the magnetic bead-template DNA complex, was removed. The supernatant was treated with an equal volume phenol: chloroform: isoamyl alcohol (25:24:1) solution to extract proteins, and then the soluble phase was precipitated with 2.6 volumes 100% ethanol. After centrifugation at 14,000 rpm for 30 min, the pellet was dissolved with nuclease-free water. *EGR1* transcripts were converted into cDNA and quantified using a OneStep RT-PCR kit (Qiagen) and a pair of primers (Table S1), visualized on native PAGE gels, and then they were quantified using Image J. For the immobilized template assay, the same procedure was followed except for collecting the pellet and supernatant fractions at the desired time points. To map TOP2B, indicated restriction enzymes were added to the reactions at the appointed time points and allowed to sit for 15 min. The pellet fraction was dissolved in 0.1 M Buffer D1. The pellet and supernatant fractions were visualized in PAGE gel (for DNA) and in SDS-PAGE, followed by silver-staining (for proteins, silver nitrate purchased from Sigma) and Western blotting (specific proteins of interest). Buffers used for transcription and immobilized template assays included freshly added protease inhibitors, 1 mM benzamidine, 0.25 mM PMSF, aprotinin (1:1000), and 1 mM Na-metabisulfite.

### Western blot and immunoprecipitation

Primary antibodies for probing phosphorylated S2 Pol II (ab5095) were obtained from Abcam (ab5095). The primary antibodies obtained for Bethyl Laboratories were as follows: BRCA1 (A300-000A), BARD1 (A300-263A), phosphorylated BRCA1 at S1524 (A300-001A), MED23 (A300-425A), and TOP2B (A300-949A). The antibodies for *α*-Tubulin (sc-8035), TOP2B (sc-25330), TFIID (sc-421), TFIIF (sc-37430), TFIIE*α* (sc-133065), CDK9 (sc-13130, and Ubiquitin (sc-8017) were from Santa Cruz Biotechnology. ELK1 (#91825) and Pol II (#2629) were purchased from Cell Signaling Technology. Each antibody was diluted in blocking solution within the range 1:500–1:3000, following to the manufacturer’s suggestion and empirical outcome. Rabbit and mouse secondary antibodies and luminol reagents for Western blotting (sc-2048) were purchased from Santa Cruz Biotechnology and diluted to 1:2000 in blocking solution for usage. For Western blotting, HEK293 lysates were collected using RIPA buffer (Cell Signaling Technology), and the protein concentrations of the lysates were quantified through Bradford assays (Bio-Rad) before SDS PAGE to compare the samples in equal amounts of total proteins. For immunoprecipitation, protein A agarose beads (Cat. # 20333, Pierce) were equilibrated with 0.15 M HEGN (50 mM HEPES, pH 7.6, 0.15 M KCl, 0.1 mM EDTA, 10% glycerol, 0.02% NP40), and TOP2B antibodies (A300-949A, Bethyl) or control IgG (sc-69876, Santa Cruz Biotechnology) were bound to the protein A beads. The antibody-beads complex was washed with a 30-bead volume 0.5 M KCl-HEGN three times each and then with same volume 0.15 M HEGN twice. The TOP2B antibody-bound protein A beads were incubated with 5 mg HeLa NE for 3 h at 4°C. The WT, SA, and SD BRCA1 proteins and control with a protein storage buffer only were added along with HeLa NE. The bead-protein complexes were washed with a 33-bead volume of 0.25 M KCl-HEGN three times. All buffer solutions included fresh protease inhibitors described above. The resultant pellet fractions were stored and subjected to silver staining and Western blotting analyses.

### Chromatin immunoprecipitation & qPCR

The ChIP experiment was conducted following the Abcam X-ChIP protocol, with mild modifications. Cell lysis buffer included 5 mM PIPES (pH 8.0), 85 mM KCl, 0.5% NP-40. Nuclei lysis buffer, including 50 mM Tris-HCl (pH 8.0), 10 mM EDTA, and 1% SDS, was added before sonication. Sonication was performed on ice at 25% amplitude for 30 s at 2 min intervals (Vibra-Cell Processor VCX130, Sonics) and was optimized to produce DNA segments ranging between +100 and +1,000 bp on a DNA gel. The cell and nuclei lysis buffers included the fresh protease inhibitors described above. The antibodies used in immunoprecipitation were Pol II (ab817, Abcam; #2629, Cell signaling, USA; A304-405A, Bethyl Laboratories, USA), phosphorylated S2 Pol II (ab5095, Abcam, USA), BARD1 (A300-263A, Bethyl Laboratories, USA), TOP2B (A300-949A, Bethyl Laboratories, USA; sc-25330, Santa Cruz Biotechnology, USA), BRCA1 (A300-000A, Bethyl Laboratories, USA), and phosphorylated BRCA1 (S1524) (A300-001A, Bethyl Laboratories, USA; NB100-200, Novus Biologicals, USA). After IP and reverse cross-linking, the DNA was purified with a Qiagen PCR purification kit. The input DNAs were quantified for qPCR analyses by nanodrop to normalize the amount of template DNAs. qPCR was performed with Platinum Tag DNA Polymerase High Fidelity (Invitrogen) under thermal cycling for 2 min at 94 degree followed by 30 cycles of 20 s at 94°C, 30 s at 55°C, and 30 s at 68°C. The resulting PCR products were analyzed through native <u>p</u>olyacryl<u>a</u>mide <u>g</u>el <u>e</u>lectrophoresis (PAGE) and then quantified with Image J. The ChIP products were also analyzed using real time PCR using SYBR Green Realtime PCR Master Mix (Toyobo) and the primers listed in Table 1 under thermal cycling as 1 min at 95°C followed by 45 cycles of 15 s at 95°C, 15 s at 55°C, and 45 s at 72°C.

### Immunofluorescence

The HeLa cells were grown on a cover-glass and were cultured for 24 h in serum-containing medium. The cells were treated with etoposide 5 μM for 1 h. For immunofluorescence analyses, the cells were fixed with 4% para-formaldehyde for 20 min and washed twice with PBS. Then the cells were permeabilized with 0.1% Triton X-100 in PBS. After blocking with 5% bovine serum albumin with PBS, the cells were incubated with anti-TOP2B (H-8, Santa Cruz, Dallas, TX) and anti-phospho-BRCA1 (Ser1524) (#9009, Cell Signaling Technology, Danvers, MA) antibodies for 3 h and secondary antibodies (anti-mouse or anti-rabbit) conjugated with Alexa Fluor 488/594 for 1 h. The nuclei were further stained with DAPI (blue, nuclear staining). Then the fluorescence images were captured using an LSM 700 laser scanning confocal microscope with an objective C-Apochromat 40×/1.2 W Corr UV-VIS-IR M27 (Carl Zeiss, Jena, Germany). Fluorophores were visualized using the following filter sets: 488-nm excitation and band-pass 420– 550 emission filter for Alexa 488; 555-nm excitation and long-pass 560 filter for Alexa 594. DAPI was visualized using 405 nm excitation and 410 nm emission long pass filters. The scale bar on the bottom right of each image indicates 5 µm.

### Chromatin immunoprecipitation & sequencing

The cell culture, serum-treatment, and ChIP experimentation were conducted as described above. Briefly, 1.5 million HEK293 cells were immunoprecipitated using an antibody, TOP2B (sc-25330, Santa Cruz Biotechnology, USA). Illumina libraries were constructed using Swift Science’s Accel-NGS Library Preparation Kit for Illumina Platforms, according to the manufacturer’s directions. Samples were quantified with qPCR using the KAPA qPCR library quant kit, following the manufacturer’s protocol, and were pooled at an equal ratio before being sequenced on the HiSeq 2500 based on qPCR concentrations (http://genomecore.wi.mit.edu/index.php/NCBISubmission).

### *In vitro* ubiquitination assay

*In vitro* ubiquitination assay with recombinant TOP2B, WT/SA/SD BRCA1, and BARD1 proteins was conducted using recombinant human ubiquitin (U-100H, R&D system, USA) and HeLa NE as a source of E1 and E2 ligases in 20 mM Tris-HCl pH 7.5, 0.2 mM DTT, 5 mM MgCl_2_, 10 μM ZnCl_2_, 3 mM ATP, 1 mM BSA, and the protease inhibitors listed in *in vitro* transcription assay. One reaction included 40 μM ubiquitin and 4 μg of HeLa NE and 50 ng of WT *EGR1* template DNA when indicated. Ubiquitination was allowed for 3.5 h at 30 °C. The reaction was stopped by the addition of 8X SDS loading buffer and unmodified and ubiquitinated recombinant TOP2B was detected using His antibody (sc-8036, Santa Cruz biotechnology, USA). For E2 ubiquitin-conjugating enzyme screening, E2 enzymes were purchased (ab139472, Abcam, USA) and the experiments were performed according to the manufacturer’s recommendations. Each of eleven E2 enzymes of UBCH1, UBCH2, UBCH3, UBCH5a, UBCH5b, UBCH5c, UBCH6, UBCH7, UBCH8, UBCH10, and UBCH13 was added to 2.5 μM. The buffer (10X) including 200 mM Tris-HCL, 2 mM DTT, 100 μM ZnCl_2_, and 10 mg/mL BSA was purified using 0.45 μm filter (Corning, USA). For UBCH2, the buffer provided with the enzymes (ab139472, Abcam, USA) was used for its DTT sensitivity. E1 enzyme provided with the kit (ab139472, Abcam, USA) and ubiquitin (U-100H, R&D system, USA) were added to 100 nM and 2 μM, respectively. Approximately, 50 nM of the recombinant SA BRCA1-BARD1 complex as an E3 ligase was added to each reaction. Recombinant full-length hTOP2B as a target protein was added at 133 nM. The ubiquitination reaction was initiated by adding Mg^2+^/ATP (ab139472, Abcam, USA), was allowed for 4 h at 37 °C before the addition of 8X non-reducing SDS gel loading buffer (without β-mercaptoethanol), and was subjected to SDS-PAGE and immunoblotting.

### Electrophoretic mobility shift assay (EMSA)

Full-length hTOP2B was diluted to targeted concentrations using TOP2B dilution buffer (10 mM HEPES, 130 mM KCl, 7.5 mM MgCl_2_, 5% glycerol, 10 μM ZnCl_2_, 0.1% NP40). For each reaction, TOP2B was incubated with 100 ng DNA in TOP2B-DNA binding buffer (13 mM HEPES, 98 mM KCl, 65 μM EDTA, 3.3% glycerol, 6.5 μM ZnCl_2_, 3.26 mM MgCl_2_, 0.05% NP40) for 40 min at room temperature. A 5% native polyacrylamide gel was made using TBE buffer and pre-run in 0.5 X TB buffer for > 30 min. Before loading onto the gel, 50% glycerol was added to each sample to 13% as the final concentration. The gel was silver-stained (Cat. 209139, Sigma-Aldrich, USA) and the band intensity was quantified using Image J. The K_d_ value with 95% confidence intervals was calculated using Prism 8 (GraphPad, Inc., San Diego, CA, USA).

### Statistical analysis

One- or two-way ANOVA was used to determine significance (*P* < 0.05) for ChIP-qPCR and RT-PCR. P-values and graphs were calculated and drawn using Prism 8 (GraphPad, Inc., San Diego, CA, USA).

### Bioinformatics

BRCA1 (GSM997540) and TOP2B (GSM2442946) ChIP-seq data were downloaded from the gene expression omnibus (GEO) database (Dellino et al., 2019; Gardini et al., 2014). The raw data in the FASTQ format were processed using the Octopus-Toolkit (version 2.1.3)(Kim et al., 2018). Briefly, the sequenced reads were trimmed using Trimmomatic (version 0.36)(Bolger et al., 2014), and then were aligned to the reference genome (hg38 assembly) using HISAT2 (version 2.1.0)(Kim et al., 2015). BRCA1 and TOP2B binding sites (peaks) were identified using HOMER (version 4.10.1)(Heinz et al., 2010) according to the following parameters: −region -size 1000 -minDist 2500. Integrative Genomics Viewer (version 2.3.69)(Robinson et al., 2011) was used to capture snapshots of the given loci with the BRCA1 or TOP2B ChIP-seq data. Heatmaps were generated for the promoter regions of the given genes using Deeptools (version 2.0)(Ramirez et al., 2016).

## Results

We previously found that DDR is coupled with productive transcriptional elongation in stress-inducible genes including *HSP70* and IEGs and that the DDR can be attributed, at least in part, to the catalytic activity of TOP2B (Bunch, 2017; Bunch et al., 2015). Therefore, we hypothesized that Top-DSB occur due to transcriptional activation, and the DDR is a manifestation of the effort to repair the DSB. In spite of the involvement of the DNA-PK and KU proteins, it is not clear which pathway among HDR, NHEJ, and precise-NHEJ is the prime repair mechanism for the Top-DSB. Notably, recent studies have also reported that DSB can be repaired by HDR in any cell cycle phase including, G_1_ (Keskin et al., 2016; McDevitt et al., 2018; Wei et al., 2015), suggesting possible competition between the two repair pathways for the Top-DSB lesion in transcription.

To understand how Top-DSB generation is regulated during transcriptional activation, we initially investigated whether BRCA1 is involved for the expression of stress-inducible genes in HEK293. Following a previously established method (Bunch et al., 2019; Bunch et al., 2015), cells were synchronized to the G_0_ phase of the cell cycle by serum starvation (S0) and then were allowed to progress into the G_1_ phase by serum supplement for 15 min (S15; **Fig. 1a**). We examined the occupancy of BRCA1 for representative hIEGs, including *FOS, MYC*, and *EGR1*, using ChIP analyses in HEK293 and MCF7 cells. BRCA1 was detected in the TSSs of the tested genes, but its occupancy was not increased upon serum induction. Then we checked for the possible activation by the phosphorylation of BRCA1 S1524 (pBRCA1), a signature modification, following DSB. DNA damage and DSB promptly induce the formation of pBRCA1, mediated by ATM and <u>a</u>taxia telangiectasia and <u>R</u>ad3-related (ATR) proteins (Cortez et al., 1999; Foray et al., 2003; Tibbetts et al., 2000). Therefore, the level of pBRCA1 was probed before and after the serum-induced transcriptional activation. The experiments showed an increase in pBRCA1, without a change in total BRCA1 occupancy, suggesting DNA damage-mediated activation of BRCA1 during processive transcription (**Fig. 1b, c**). In addition, ChIP-qPCR showed that BRCA1 KD clearly decreased the level of total Pol II and <u>s</u>erine <u>2</u>-phosphorylated CTD of <u>Pol II</u> (S2 Pol II), and TOP2B in the *EGR1* gene body, indicating reduced transcriptional activity in the gene in spite of the serum induction in the absence of BRCA1 (**Fig. 1d, e**).

**Fig. 1.**
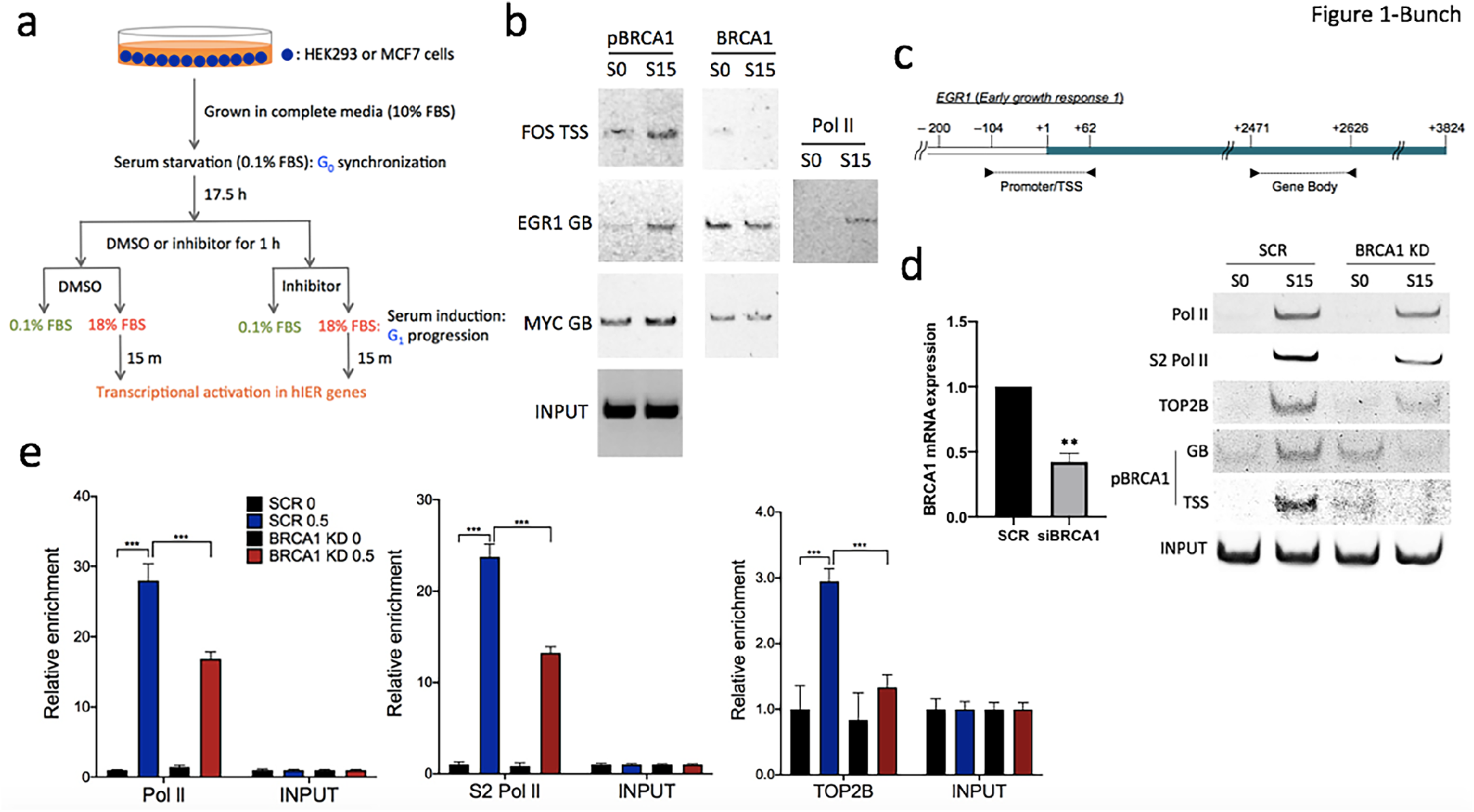
BRCA1 regulates serum-induced transcriptional activation. (**a**) Schematic overview of cell cycle synchronization and serum-induced transcriptional activation in hIEGs with or without chemical kinase inhibitors. FBS, fetal bovine serum. (**b**) ChIP-PCR showing BRCA1 and pBRCA1 occupancy before and after transcriptional activation at *FOS, MYC*, and *EGR1*. S0, serum-starved cells at G_0_; S15, serum-induced cells. (**c**) Schematic of the *EGR1* genomic region. Primers to amplify TSS and gene body in ChIP-qPCR are marked as black arrows. (**d**) Left, BRCA1 mRNA expression in scrambled siRNA (SCR) control vs BRCA1 KD cells. Error bars show standard deviations (s.d., n = 3). **P < 0.005. Right, ChIP-PCR showing reduced S2 Pol II and TOP2B recruitment and pBRCA1 formation in activated *EGR1* gene. GB, gene body. (**e**) ChIP-qPCR showing impaired Pol II, S2 Pol II, and TOP2B recruitment upon gene activation at *EGR1* in BRCA1 KD cells. Input chromatin was amplified using a *EGR1* GB primer set and used as normalizer. Error bars show s.d. (*n* = 4). ***P < 0.005.

BRCA1 KD using siRNA species prevented its expression at the mRNA and protein levels (**Fig. 2a**). We also examined BARD1 because this protein can form a heterodimer with BRCA1, a complex which constitutes a functional unit in the cell (Baer and Ludwig, 2002). BARD1 KD decreased the protein levels of not only BARD1 but also BRCA1 (**Fig. 2a**), supporting a mutually interdependent requirement for stability between the two proteins (Choudhury et al., 2004). Top-DSB occurs in serum-inducible, immediate early genes including *EGR*1, and the catalytic activity of TOP2B and DDR signaling are required for the processive transcription in these genes (Bunch et al., 2015). Either BARD1 or BRCA1 KD caused the mRNA expression of *EGR1*, a representative hIEG, to be lower than the SCR control when the gene expression was induced by serum (**Fig. 2b**), suggesting that these proteins are important for *EGR1* transcription.

**Fig. 2.**
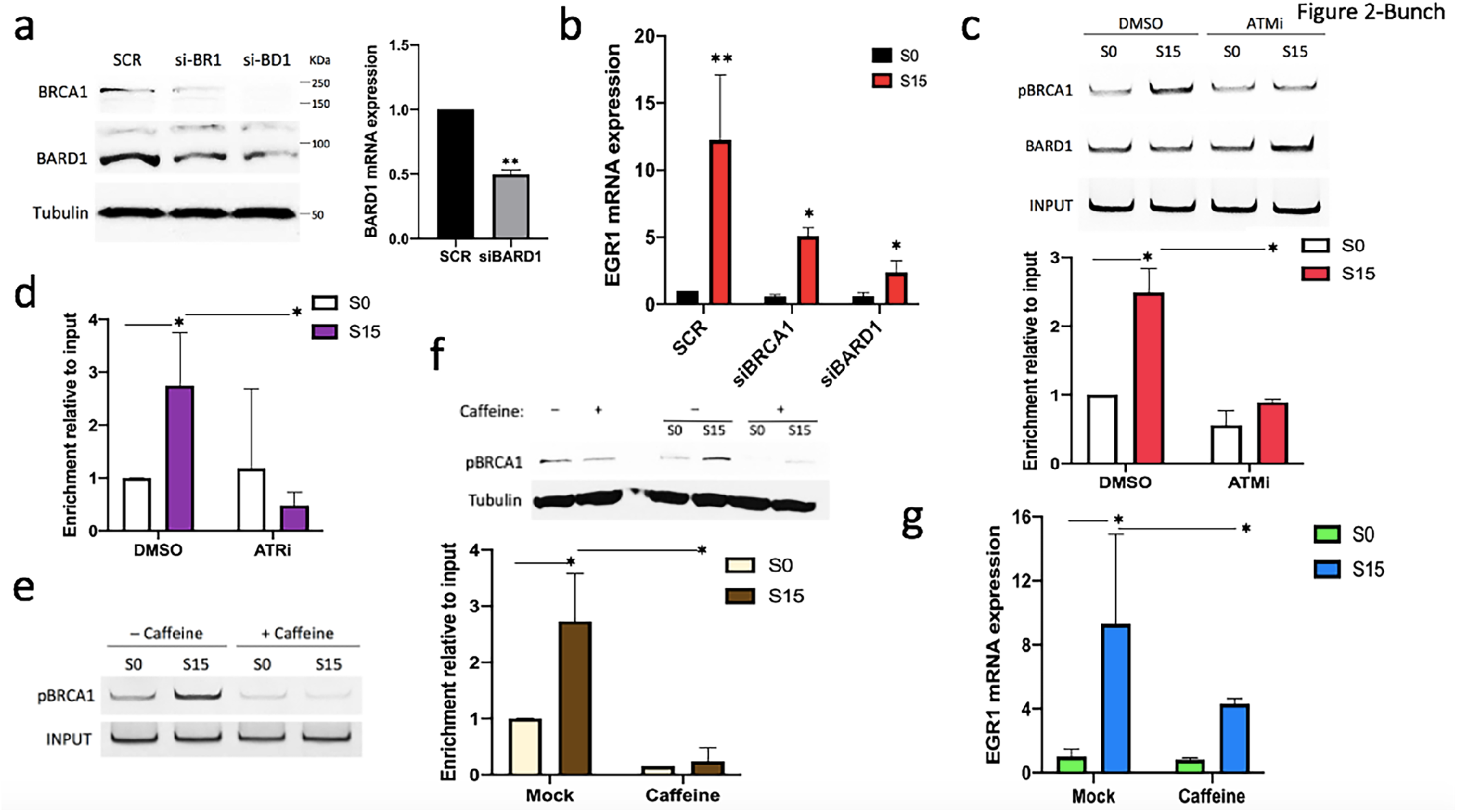
BRCA1 is phosphorylated by ATM and ATR upon transcriptional activation. (**a**) BRCA1 and BARD1 KD validation. Left, immunoblotting data showing the protein level of BRCA1, BARD1, and *α*-Tubulin (Tubulin, loading control, 70 µg cell lysate/lane) using siRNA species targeting BRCA1 (si-BR1) or BARD1 (si-BD1). Right, RT-qPCR data showing the mRNA level of BARD1. GAPDH was used as a normalizer. Error bars show in s.d. (*n* = 3). **P < 0.005. (**b**) RT-qPCR data showing the effects of BRCA1 or BARD1 KD on *EGR1* mRNA expression. GAPDH was used as a reference gene. Error bars show s.d. (*n* = 3). *P < 0.05 and **P <0.01. (**c**) Upper, ChIP-PCR showing that the occupancy of pBRCA1 in the TSS of *EGR1* is markedly lower in the presence of KU55933 (ATMi) during transcriptional activation than the DMSO-treated control (DMSO). Bottom, ChIP-qPCR of pBRCA1 with or without ATMi in *EGR1* TSS. Error bars show s.d. (*n* = 2). *P < 0.05. (**d**) ChIP-qPCR of pBRCA1 with or without VE-821 (ATRi) in *EGR1* TSS. Error bars show s.d. (*n* = 2). *P < 0.05. (**e**) Left, ChIP-PCR showing that caffeine, which inhibits ATM and ATR, alleviates the accumulation of pBRCA1 on *EGR1* TSS in transcriptional activation. Right, ChIP-qPCR of pBRCA1 on *EGR1* TSS. Error bars show s.d. (*n* = 3). *P < 0.05. (**f**) Immunoblotting showing that caffeine reduces pBRCA1 in cells and upon transcriptional activation. α-Tubulin was used as a loading control. (**g**) qRT-PCR showing the reduction of *EGR1* mRNA level in caffeine-treated cells upon transcriptional activation. *β-ACTIN* as a reference and normalizer. Error bars show in s.d. (*n* = 3). *P < 0.05.

To understand the function of BRCA1 phosphorylation at S1524, small molecule inhibitors of ATM and ATR, KU55933 and VE821, were added for 1.25 h before and during the serum induction, respectively (**Fig. 1a**). Our ChIP-PCR data showed reduced levels for pBRCA1 during exposure to these inhibitors in S15 samples in the TSS of *EGR1*, although the occupancy of BARD1 was not altered (**Fig. 2c, d**). Consistent with these findings, caffeine, a compound that inhibits the kinase activities of both ATM and ATR (Sarkaria et al., 1999) interfered with pBRCA1 accumulation at the *EGR1* gene upon transcriptional activation (**Fig. 2e**). In addition, caffeine exposure reduced the level of pBRCA1 both with and without cell cycle synchronization, confirming that these kinases are indeed responsible for BRCA1 phosphorylation during transcriptional activation (**Fig. 2f**). Consistently, caffeine treatment reduced the *EGR1* mRNA level in S15 samples (**Fig. 2g**). These data demonstrate that BRCA1 is phosphorylated by ATM and ATR during transcriptional activation and suggest that pBRCA1 is important for gene expression of *EGR1*.

We validated the functional importance of BRCA1 phosphorylation at S1524 for transcriptional activation using *in vitro* transcription assay. For this purpose, recombinant WT BRCA1 (220 KDa) and two BRCA1 S1524 mutants, S1524A (phospho-null, SA) and S1524D (phospho-mimic, SD) were cloned and purified from *E. coli* and validated by immunoblotting (**Supplementary Fig. 1A–C**). The resulting WT, SA, and SD BRCA1 proteins were compared using the *in vitro* transcription assay established in our previous study (Bunch et al., 2014). In brief, transcriptional PIC was formed on a biotinylated *EGR1* template DNA construct that included the promoter and TSS (–423 to +323, **Supplementary Fig. 2A**) using HeLa NE (**Supplementary Fig. 2B**). For the *in vitro* biochemical analyses, including immobilized template and transcription assays, we utilized HeLa NE because the conditions and methods have been well-established and validated in the studies of others and in our previous investigations (**Fig. 3a**)(Bunch et al., 2014; Kim et al., 1998; Lin and Carey, 2012). PIC formation on the *EGR1* TSS template was confirmed by probing Pol II, general transcription factors including TFIID, CDK9, and MED23, and ELK1, a promoter-binding transcriptional factor specific to the *EGR1* gene (Shan et al., 2014), using immunoblotting (**Fig. 3b**). WT and mutant BRCA1 proteins were added along with NE only (T1) or both with NE and 3–5 min after NTP (T2; **Fig. 3a**). We hypothesized that the recombinant BRCA1 proteins supplemented at T1 could affect PIC formation, while the proteins at T2 could influence both PIC formation and transcriptional initiation and promoter-proximal pausing. As an alternative to the use of radioactive rNTP and sequencing gel electrophoresis, *EGR1* transcripts from each experimental condition were converted into complementary DNA and quantified by PCR, using a pair of *EGR1*-specific oligonucleotides, amplifying from +1 to +332 (Table S1). Both T1 and T2 addition of the SD BRCA1 stimulated transcription, relative to WT and SA and yet the effect was more noticeable at T2 (**Fig. 3c**). In addition, to exclude the preexisting *EGR1* transcripts in NE from being included in the quantification (**Supplementary Fig. 2C**), we modified 14 nucleotides from the template DNA between +140 and +160 to generate an *EGR1* transcript (**Supplementary Fig. 3A**) that would be distinguished from the native one (**Supplementary Fig. 3B**). When the nascent transcripts from mod-EGR1 were compared among the recombinant BRCA1 species at T2, the results confirmed that phosphomimetic SD BRCA1 enhances transcription, but WT or SA does not (**Fig. 3c**; **Supplementary Fig. 3C**). SD showed the most transcriptional activation of *in vitro* transcription, by approximately over 2.5 fold compared to the WT control (**Fig. 3d**). Together with the cell-based data (**Fig. 1, 2**), these results strongly indicate that phosphorylation of BRCA1 at S1524 is important for the active transcription of the *EGR1* gene.

**Fig. 3.**
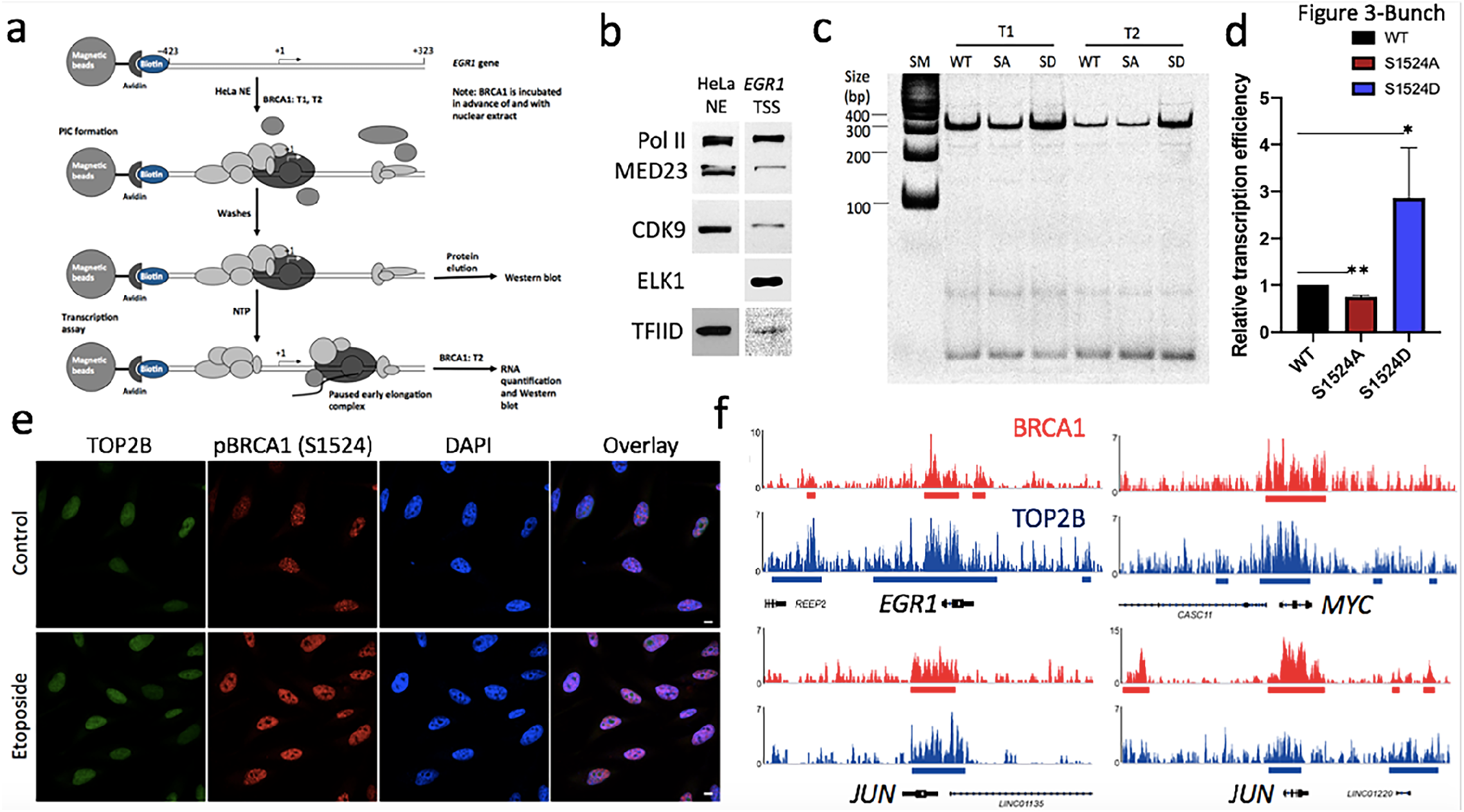
BRCA1 phosphorylation at S1524 is important for transcriptional activation. (**a**) Schematic overview of immobilized template and *in vitro* transcription assays. Recombinant BRCA1 proteins were added to the reaction before and during PIC formation (T1) and both during PIC formation and after transcription initiation (T2). (**b**) Validation of PIC formation on EGR1 TSS (–432 to +332) using immunoblotting. (**c**) In vitro transcription assay using recombinant WT, S1524A (SA), and S1524D (SD) BRCA1 showing that SD activates *EGR1* transcription, compared to WT and SA. (**d**) Quantification of the efficiency of S1524A and S1524D BRCA1 mutant proteins in stimulating transcription, relative to that of WT BRCA1. Error bars show s. d. (*n* = 3). *P < 0.02; **P <0.002. (**e**) Immunofluorescence-confocal microscopy results showing increased levels of TOP2B and pBRCA1 in nuclei upon etoposide treatment. (**f**) Chromosome views of total BRCA1 (red) and TOP2B (etoposide-trapped, blue) on representative hIEGs, *EGR1, JUN, MYC*, and *FOS*, illuminating genomic colocalization and functional collaboration.

Although BRCA1 is involved in the transcription of a diverse group of genes, the mechanisms through which it functions in transcription are incompletely understood (Mullan et al., 2006; Welcsh et al., 2002). Previous studies have shown that BRCA1 phosphorylation at S1524 is induced by DNA damage (Cortez et al., 1999) and that the representative hIEGs listed above require DNA breaks mediated by TOP2B for Pol II pause release and active Pol II elongation (Bunch et al., 2015; Madabhushi et al., 2015). Notably, a recent study found that BRCA1 regulates the resolution of TOP2B-DNA adducts during the transcriptional activation of estrogen receptor-activated genes in the G_1_ phase (Sasanuma et al., 2018). Another study showed that the function of TOP2A, which is highly homologous to TOP2B except in its C-terminal domain (Linka et al., 2007) is regulated by the ubiquitin-ligase function of the BRCA1-BARD1 complex during S phase (Lou et al., 2005). We attempted to examine the relation between TOP2B and pBRCA1. It was asked whether TOP2B-DNA adducts induced by etoposide increase BRCA1 phosphorylation at S1524 by treating the cells with etoposide at 5 µM for 1 h. The data show that pBRCA1 increased for TOP2B-DNA adducts in the nucleus (**Fig. 3e**; **Supplementary Fig. 4A**), a finding which suggests that DSBs caused by the abortive catalysis of TOP2B lead to the increase in pBRCA1. In addition, we examined specific locations of BRCA1 and TOP2B in hIEGs using the ChIP-seq data (GSM2442946 and GSM997540)(Dellino et al., 2019; Gardini et al., 2014). The colocalization pattern of BRCA1 and TOP2B was evident when the factors were shown in chromosome viewers for the representative hIEGs, including *EGR1, JUN, MYC*, and *FOS* (**Fig. 3f**). When we examined TOP2B trapped by etoposide in the protein-coding genes occupied by BRCA1, these proteins colocalized in a large number of the genes (*n* = 13,903; **Fig. 4**). Their colocalization trend was evident and almost identical also when the occupancy of BRCA1 was visualized in the protein-coding genes occupied by etoposide-trapped TOP2B (**Supplementary Fig. 4B**). These results suggested that TOP2B and BRCA1 could modulate gene expression by regulating transcription.

**Fig. 4.**
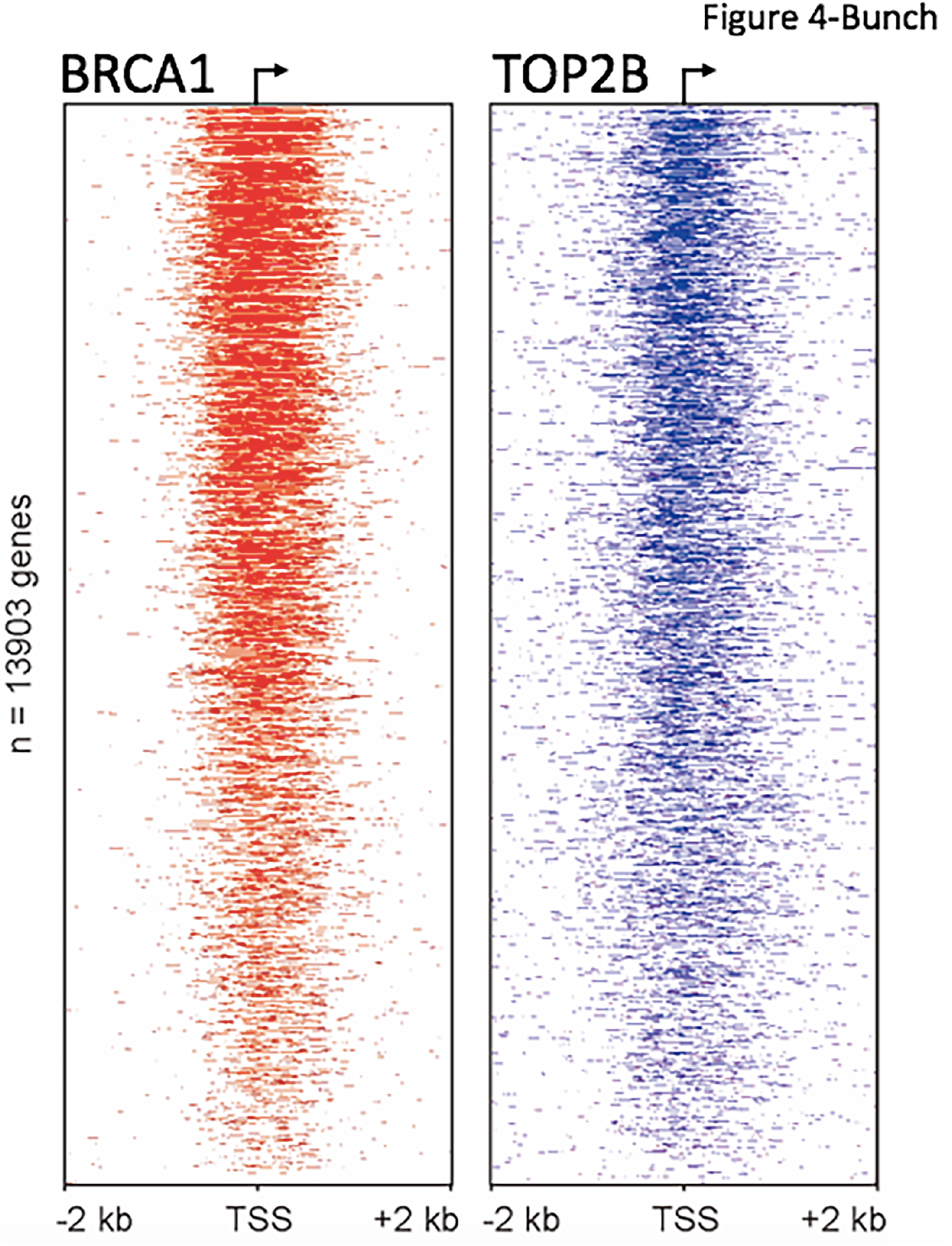
TOP2B active sites and BRCA1 binding sites are overlapped genome-wide. Heat maps of TOP2B and BRCA1 sorted by BRCA1 gene occupancy (*n* = 13,903 protein coding genes only). Normalized, input-adjusted total BRCA1 (GSM2442946) and TOP2B (GSM997540) ChIP-seq reads.

Therefore, we next hypothesized that the BRCA1-BARD1 complex might be involved in TOP2B regulation. To test this hypothesis, we first sought to determine whether TOP2B and BRCA1-BARD1 complex could physically interact using IP against NE, followed by Western blotting. The control experiment with IgG did not pull down the proteins of interest (**Fig. 5a**). The results using specific antibodies showed little recognizable physical interaction between BRCA1 and TOP2B proteins (**Fig. 5a**). On the other hand, we found a strong and stable interaction between TOP2B and BARD1 that was sustained through a series of stringent high-salt washes (**Fig. 5a**). Intriguingly, recombinant WT BRCA1, added to the IP along with NE, appeared to sequester BARD1 from TOP2B, interfering with their interaction (**Fig. 5a**). By contrast, the SA and SD BRCA1 proteins did not interfere with the interaction between BARD1 and TOP2B, which was sustained almost to a level comparable to that of the control without WT BRCA1 supplement (**Fig. 5a**). In addition, when recombinant WT BRCA1 was supplemented in the reaction, BARD1 dissociated from TOP2B, resulting in reduced ubiquitination signals (**Fig. 5a**). By contrast, neither the SA nor the SD BRCA1 proteins interfered with the BARD1-TOP2B interaction, and ubiquitination signals were observed in both samples, although to different degrees (**Fig. 5a**). The ubiquitination signal was stronger with SA BRCA1 (**Fig. 5a**). These results suggested that BRCA1 regulates the physical interaction of BARD1 with TOP2B and that the phosphorylation of the residue S1524 in BRCA1 may be important in regulating the BARD1-TOP2B interaction required for TOP2B ubiquitination.

**Fig. 5.**
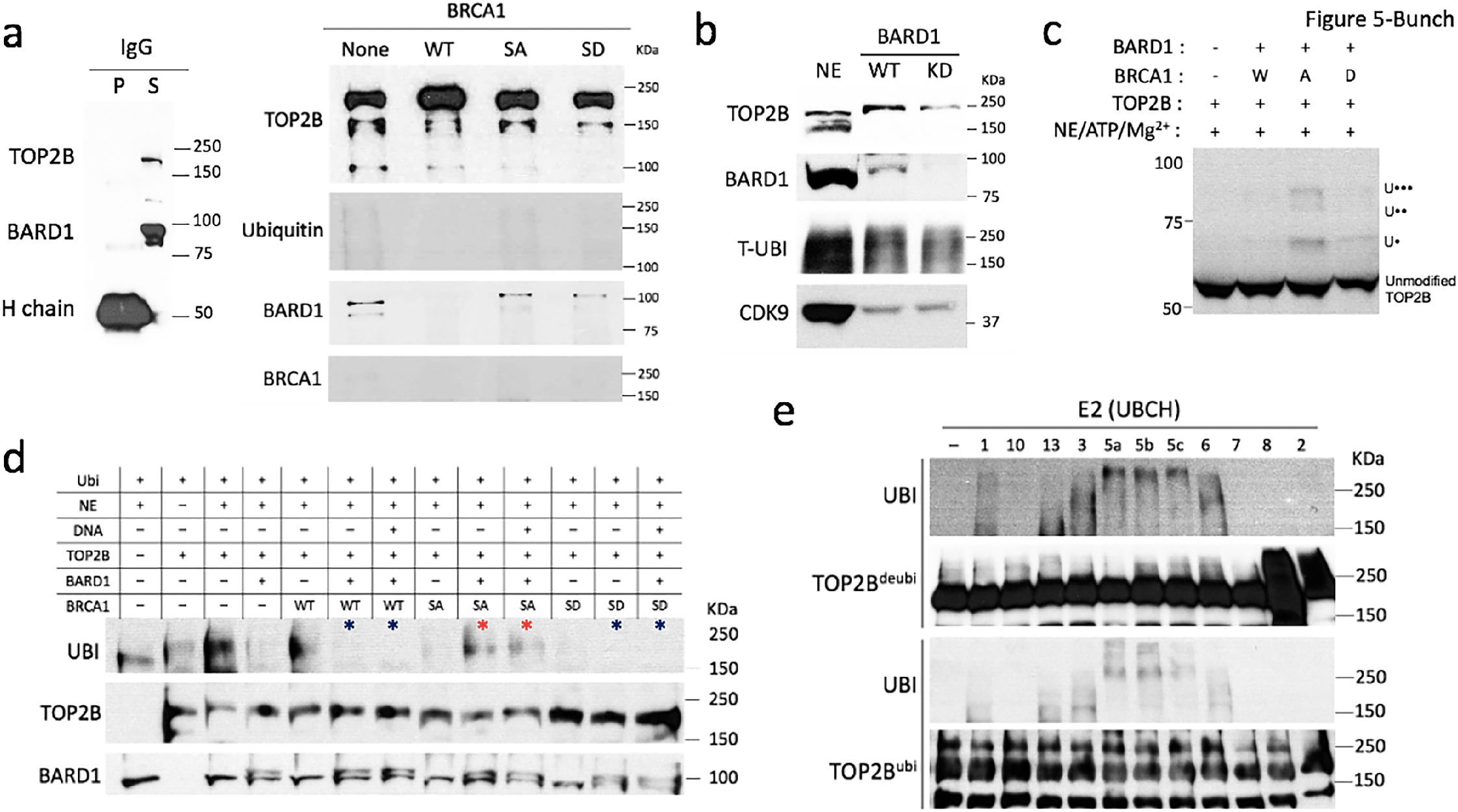
BRCA1-BARD1 binds to and ubiquitinates TOP2B in BRCA1 S1524 phosphorylation-dependent manner. (**a**) Left, immunoprecipitation with control <u>i</u>mmuno<u>g</u>lobulin <u>G</u> (IgG) against 5 mg of HeLa NE. P, pellet (bound) and S, supernatant (unbound) fraction. H chain, IgG heavy chain. Right, immunoprecipitation of TOP2B antibody against NE, followed by immunoblotting indicating a stable interaction between TOP2B and BARD1. BRCA1 did not appear to strongly bind toTOP2B. The level of BARD1 associated with TOP2B appeared positively correlated with the degree of suspected TOP2B ubiquitination. (**b**) Immunoblotting data showing that BARD1 KD decreases the level of TOP2B proteins and ubiquitination in the nuclei. CDK9 was used as a loading control. NE was included as a technical control. (**c**) *In vitro* ubiquitination assay followed by immunoblotting showing discrete bands at about 56, 68, 81, and 91 KDa for 0, 1 (U•), 2 (U• •), and 3 (U• • •) ubiquitin proteins ligated to TOP2B^1-566^ by SA BRCA1-BARD1. W, WT BRCA1; A, SA BRCA1; D, SD BRCA1. (**d**) *In vitro* ubiquitination assay and immunoblotting showing the ubiquitination of FL-TOP2B (TOP2B) by SA BRCA1-BARD1. Blue and red asterisks for comparison of WT, SA, SD BRCA1, together with BARD1, regarding their effectiveness as an E3 ligase to TOP2B. (**e**) In vitro ubiquitination assay and immunoblotting showing the E2 enzymes, UBCH1, UBCH3, UBCH5a, UBCH5b, UBCH5c, UBCH6, and UBCH13, that cooperate with the BRCA1-BARD1 complex to ubiquitinate hTOP2B-FL, TOP2B^ubi^ and TOP2B^deubi^.

To further investigate the relationship between TOP2B and BARD1, the levels of TOP2B protein were examined, comparing the nuclear extracts from WT and BARD1 KD cells, using immunoblotting. The data showed a notable reduction of TOP2B in BARD1 KD cells, which suggested TOP2B stability to be dependent on BARD1 (**Fig. 5b**; **Supplementary Fig. 5A**). In addition, the total ubiquitination signals between proteins with electrophoretic mobilities corresponding to 150 and 250 KDa were mildly reduced in the BARD1 KD cells (**Fig. 5b**). Although the source of the ubiquitin signals cannot be distinguished from this experiment, and it is unclear which proteins are modified, ubiquitination became reduced in a manner dependent on the protein levels of BARD1 and TOP2B (**Fig. 5a, b**; **Supplementary Fig. 5A**). We interpreted these findings to indicate either: (1) cellular TOP2B is present in an ubiquitinated form, so that TOP2B protein reduction itself results in the decreased ubiquitin signals. Alternatively, (2) ubiquitination may regulate TOP2B stability so that the reduced ubiquitination leads to a decline in protein level in BARD1 KD cells.

We were then prompted to ask whether the BRCA1-BARD1 complex ubiquitinates TOP2B directly, by acting as an E3 ligase and to identify the role of BRCA1 phosphorylation in TOP2B ubiquitination through *in vitro* ubiquitination analyses. Recombinant TOP2B containing 1–566 amino acids (TOP2B^1-566^) and full-length BARD1 were purified from bacteria and verified by immunoblotting (**Supplementary Fig. 5B, C**; **Table S1**). To supply unidentified E1 and E2 enzymes, a small amount of NE was utilized. His^6^-tagged TOP2B^1–566^ was incubated with WT, SA, or SD BRCA1 and BARD1 in the in *vitro* ubiquitination and the results were analyzed by immunoblotting using an anti-His antibody. The controls, WT BRCA1-BARD1 complex or BARD1 only without TOP2B^1–566^ did not display the shifted band characteristic of ubiquitinated TOP2B^1–566^ (**Supplementary Fig. 5D**). Importantly, SA BRCA1 and BARD1 resulted in formation of mono-, di-, and tri-ubiquitinated TOP2B^1–566^, calculated from the log_10_MW plot of molecular migration (**Fig. 5c**; **Supplementary Fig. 5E**). The WT and SD BRCA1 incubation did not result in significant levels of ubiquitinated TOP2B^1–566^. These results showed that the BRCA1-BARD1 complex ubiquitinates TOP2B, and this modification is dependent on the phosphorylation status of BRCA1 at S1524, as unphosphorylatable BRCA1 S1524 supports discrete ubiquitination of TOP2B 1–566 amino acids (**Fig. 5c**). In addition, a full-length TOP2B purified from human cells was tested for ubiquitination *in vitro* (**Fig. 5d**; **Supplementary Fig. 5F**). Supporting our current inferences, the purified full-length TOP2B was partially ubiquitinated and was further ubiquitinated by addition of NE (**Fig. 5d**). Recombinant BARD1 purified from bacteria (migrates slightly above endogenous BARD1, because of 50 amino acids derived from vector sequence) suppressed TOP2B ubiquitination, probably due to the lack of protein modifications in the prokaryotic cells and the competition with endogenous BARD1. However, it was shown that TOP2B was ubiquitinated by SA BRCA1-BARD1, but not rarely by WT- and SD BRCA1-BARD1 (**Fig. 5d**). Together with **Fig. 5c**, these data suggest that the unphosphorylated SA BRCA1-BARD1 is a potent E3 ligase for TOP2B.

To identify the E2 enzyme(s) that cooperates with the BRCA1-BARD1 to ubiquitinate TOP2B, eleven E2 ubiquitin-conjugating enzymes (UBCH1, UBCH2, UBCH3, UBCH5a, UBCH5b, UBCH5c, UBCH6, UBCH7, UBCH8, UBCH10, and UBCH13) were screened for their ability to ubiquitinate recombinant TOP2B *in vitro*. Because the recombinant human full-length TOP2B obtained from HEK293F cells was partially ubiquitinated (hereafter, TOP2B^ubi^), we added the deubiquitinase USP2 to wash steps during the YFP-tag purification steps. Subsequent washes and FPLC purification remove USP2 to yield de-ubiquitinated full-length TOP2B (hereafter, TOP2B^deubi^; **Supplementary Fig. 6A**). These TOP2B^ubi^ and TOP2B^deubi^ were used as target proteins for ubiquitination in the *in vitro* ubiquitination assay. The non-phosphorylatable SA BRCA1-BARD1, as an E3 enzyme, and recombinant E1 enzyme were included in each reaction. We found that UBCH3 and UBCH6 caused shifted bands between 190 and 250 KDa protein markers by modifying TOP2B^ubi^ and TOP2B^deubi^ (**Fig. 5e**). These distinguishable bands could be attributable to short ubiquitin chains or mono-ubiquitin attached to TOP2B. UBCH5a, UBCH5b, and UBCH5c generated dramatically shifted bands over 250 KDa, suggesting they catalyze poly-ubiquitination of TOP2B (**Fig. 5e**). We included small amounts of E1 and E3 enzymes but E2 and E3 enzymes can be ubiquitinated to transfer ubiquitin to their substrates. Therefore, we probed His^6^-tagged E1, E2, and E3 enzymes using an anti-His^6^ antibody to validate that the ubiquitination signals above 190 KDa were of TOP2B (**Supplementary Fig. 6B**). These results suggest that the BRCA1-BARD1 complex can interact with UBCH3, UBCH5, and UBCH6 enzymes to ubiquitinate TOP2B to different ubiquitin chain lengths, perhaps for distinctive functions or outcomes.

Next, the dynamics of TOP2B and BRCA1-BARD1 complex interaction on the *EGR1* TSS was monitored using the immobilized template assay. Initially, the binding sites of these proteins on the TSS of the *EGR1* gene were investigated. The biotinylated *EGR1* template (–432 to +323, 120 ng per reaction) was first incubated with NE to assemble the PIC, and unbound or loosely associated proteins were washed off. Then the complex was treated with or without the restriction enzyme SacII, which cuts the template at +92 (**Fig. 6a, b**). The proteins and digested DNA derived from these reactions were visualized with SDS-PAGE and silver staining, and native PAGE, respectively, confirming the proper fragmentation of the template and proteins associated with each fraction (**Fig. 6a, b**). Whereas BRCA1 was not significantly associated with the template DNA, TOP2B was found solely in the pellet fraction, along with BARD1, as observed in Western blotting. These proteins of interest did not bind to the beads only control without the *EGR1* TSS template (**Supplementary Fig. 7A**). These data indicated that TOP2B and BARD1 bind to the TSS of *EGR1* between –432 and +92 (**Fig. 5a**). NTP addition after the PIC formation on the template did not lead to significant differences in these properties (**Fig. 6a**).

**Fig. 6.**
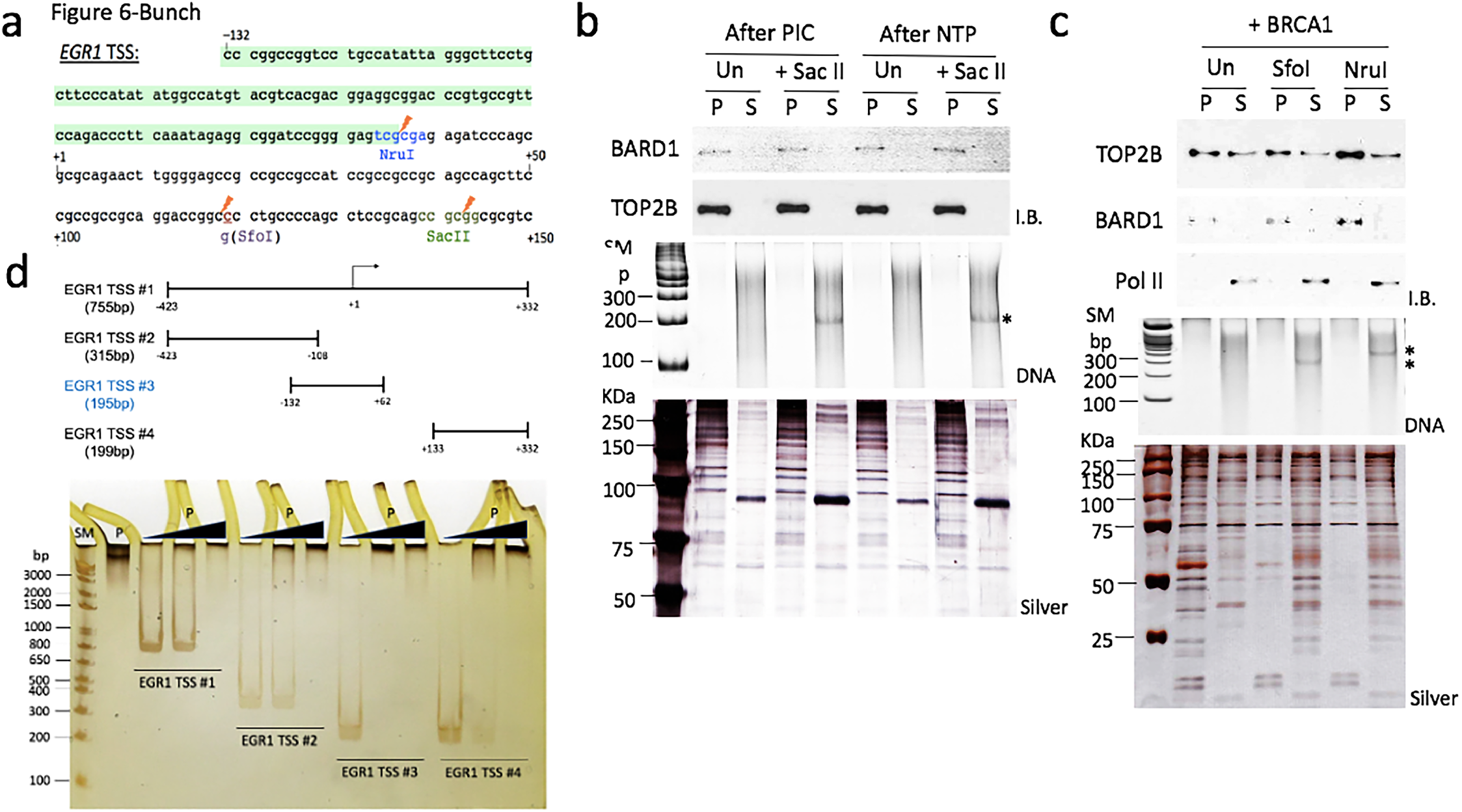
TOP2B and BARD1 bind to EGR1 TSS between –132 and –15. (**a**) DNA sequence of *EGR1* TSS (–132 to +332). Orange flash signs indicate the restriction enzyme sites for NruI, SfoI, and SacII used to map the factor binding region. Colored letters indicate those that were subjected to mutations for the purposes indicated in the text. Altered sequences are presented under the original ones. TOP2B and BARD1 mutual binding site mapped in this study was boxed with light green. (**b**) Immobilized template assay combined with restriction enzyme digestion using EGR1 *TSS* (–432 to +332) and SacII (Sac, to digest at +92), followed by immunoblotting. SacII added immediately after PIC formation or NTP addition. Undigested template DNA is marked as Un. I.B., immunoblotting probing BARD1 and TOP2B; DNA, native PAGE detecting the released DNA fragment (241 nt) after SacII digestion; Silver, silver staining visualizing the proteins bound on the DNA. P, pellet; S, supernatant fraction. (**c**) Immobilized template assay using an *EGR1* template (−132 to +332) in the presence or absence of BRCA1, combined with restriction enzyme digestion with NruI and SfoI that cut at −15 and +68, respectively. Recombinant WT BRCA1 was added at T2, immediately before the template was digested by restriction enzymes. The pellet and supernatant fractions were analyzed with immunoblotting. TOP2B was bound in the pellet fractions along with BARD1, mapping them both between –132 and –15 of *EGR1* TSS regardless of WT BRCA1 presence. (**d**) Upper panel, a diagram of the four EGR1 TSS fragments. Bottom panel, EMSA followed by silver staining showing TOP2B^ubi^ binding to *EGR1* TSS fragments with different affinities. The strongest binding was observed with *EGR1* TSS #3 (–132 to +62). SM, size marker; P, TOP2B^ubi^. The second lane shows TOP2B^ubi^ only.

The association between TOP2B and the *EGR1* TSS was monitored in the presence of BRCA1. Recombinant WT BRCA1 was supplemented at T2. The addition of WT BRCA1 increased the total amount of TOP2B associated with the *EGR1* template (–132 to +323), compared to the samples without it (**Fig. 6c**). A noticeable fraction of TOP2B was released from the template and detected in the supernatant fraction. Interestingly, BRCA1 supplementation at T2 increased levels of Pol II in the supernatant fraction (**Fig. 6c**). In addition, a SfoI site was introduced into the 25 bp upstream of +92 by a site-directed point mutation to further map the TOP2B binding site (**Fig. 6a**; **Supplementary Fig. 7B**). The proteins in the pellet and supernatant fractions and the digested DNA released from the pellet fraction are shown in silver staining and native PAGE in **Fig. 6c**. SfoI and NruI digestion, which cut the template DNA at +68 and –15, respectively, indicated that TOP2B and BARD1 still bind the regions between –132 and –15 (**Fig. 6c**; **Supplementary Fig. 7C**). Consistent with the IP results (**Fig. 5**), these experiments suggest the colocalization and physical interaction between TOP2B and BARD1 in the *EGR1* TSS. To assess the affinity between *EGR1* TSS and TOP2B, we performed electrophoretic mobility shift assay (EMSA) using TOP2B^ubi^ and different fragments of *EGR1* template DNA (**Fig. 6d**; **Supplementary Fig. 8A**) Although a large amount of TOP2B bound to all the fragments tested, the EGR1 TSS #3 (–132 to +62) bound to TOP2B^ubi^ with the strongest affinity (**Fig. 6d**).

TOP2B^deubi^ and TOP2B^ubi^ (Deu and Ubi in **Fig. 7a**, respectively) were incubated with 100 ng of EGR1 TSS #3 in increasing concentrations (up to 217 nM) and the reactions were visualized using silver staining (**Fig. 7b, c**; **Supplementary Fig. 5F, 6, 8B**,**C**). Strikingly, TOP2B^deubi^ dramatically lost its affinity for the DNA (**Fig. 7b, c**; **Supplementary Fig. 8C**). The Kd value for TOP2B^ubi^ and EGR1 TSS #3 binding is 59.9 ± 6.1 nM (**Fig. 7c**; **Supplementary Fig. 8B**) These data demonstrate that TOP2B ubiquitination enhances binding to *EGR1* TSS. To further understand the role of BRCA1 phosphorylation at S1524 in TOP2B binding to the gene, we compared WT, SA, and SD BRCA1 for TOP2B binding to the *EGR1* TSS during transcription. Recombinant BRCA1 species were added at T1 or T2, and the pellet and supernatant fraction were separated with or without NTP addition.

TOP2B was tightly bound to the *EGR1* template without NTP addition, before transcriptional initiation (**Fig. 7d**). Strikingly, SA increased the level of TOP2B stably associated with the *EGR1* template to levels that exceeded WT BRCA1 and SD BRCA1 (**Fig. 7e, f**). Subsequently, less TOP2B was released into the supernatant for the SA BRCA1 sample (**Fig. 7f**). Together with the biochemical data shown in **Fig. 5–7**, these results suggest that the residue S1524 in proximity to the <u>BR</u>CA1 <u>C</u>-terminus (BRCT) domain specialized in the binding of phosphoproteins (Liang et al., 2017) may be critical for the BRCA1-BARD1 functional interaction as an E3 ubiquitin ligase. We interpret this finding to mean that SA allows BRCA1-BARD1 to interact with and ubiquitinate TOP2B, which stabilizes TOP2B association with the DNA. The phosphomimetic SD form, on the other hand, is less capable of permitting BRCA1-BARD1-mediated TOP2B ubiquitination, which leads to TOP2B dissociation from the DNA (**Fig. 5a–e**; **7d, e**).

**Fig. 7.**
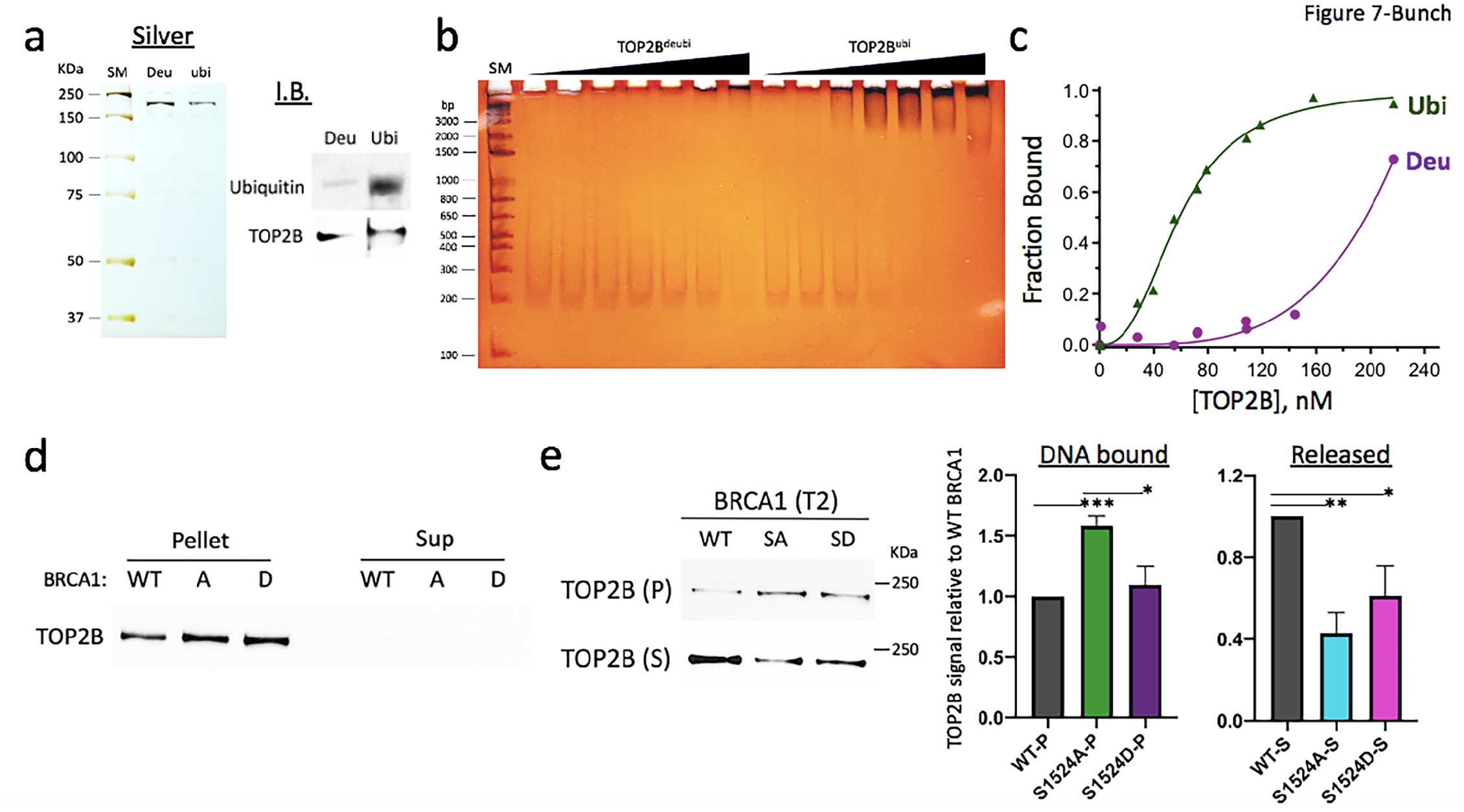
BRCA1 phosphorylation controls TOP2B ubiquitination and DNA binding affinity. (**a**) Purified TOP2B^ubi^ (ubi) and TOP2B^deubi^ (Deu) shown by silver staining and immunoblotting. (**b**) EMSA comparing TOP2B^ubi^ vs TOP2B^deubi^ for their binding affinity to EGR1 TSS #3 (–132 to +62). Silver-stained. SM, DNA size marker. (**c**) Ubiquitinated TOP2B binds to DNA with much higher affinity than deubiquitinated one. A plot summarizing EMSA to derive K_D_ values. (**d**) Immobilized template assay results. TOP2B association with *EGR1* TSS (Pellet, –432 to +332) before addition of NTP. Recombinant BRCA1 species were added at T1. (**e**) Immobilized template assay results. Left, TOP2B associated (Pellet, P) with and dissociated from EGR1 TSS (Supernatant, S) after NTP and BRCA1 protein addition at T2. Right, quantification of TOP2B in pellet (DNA bound) and in supernatant (Released) in immobilized template assays. Error bars in s.d. (*n* = 3). *P < 0.05, **P < 0.01, ***P < 0.005.

Together, cell-based, genomic, and biochemical analyses indicate a novel model in which the BRCA1-BARD1 complex is involved in the transcription of hIEGs through modulating TOP2B. In the resting state of transcription, BRCA1-BARD1 complex ubiquitinates TOP2B for a stable binding to the TSS. In transcriptional activation and pause release, Top-DSB activates ATM and ATR to phosphorylate BRCA1 at S1524. Phosphorylation of BRCA1 at S1524 controls the interaction between TOP2B and BARD1, resulting in decreased TOP2B ubiquitination, which destabilizes the TOP2B binding to the DNA (**Fig. 8**).

**Fig. 8.**
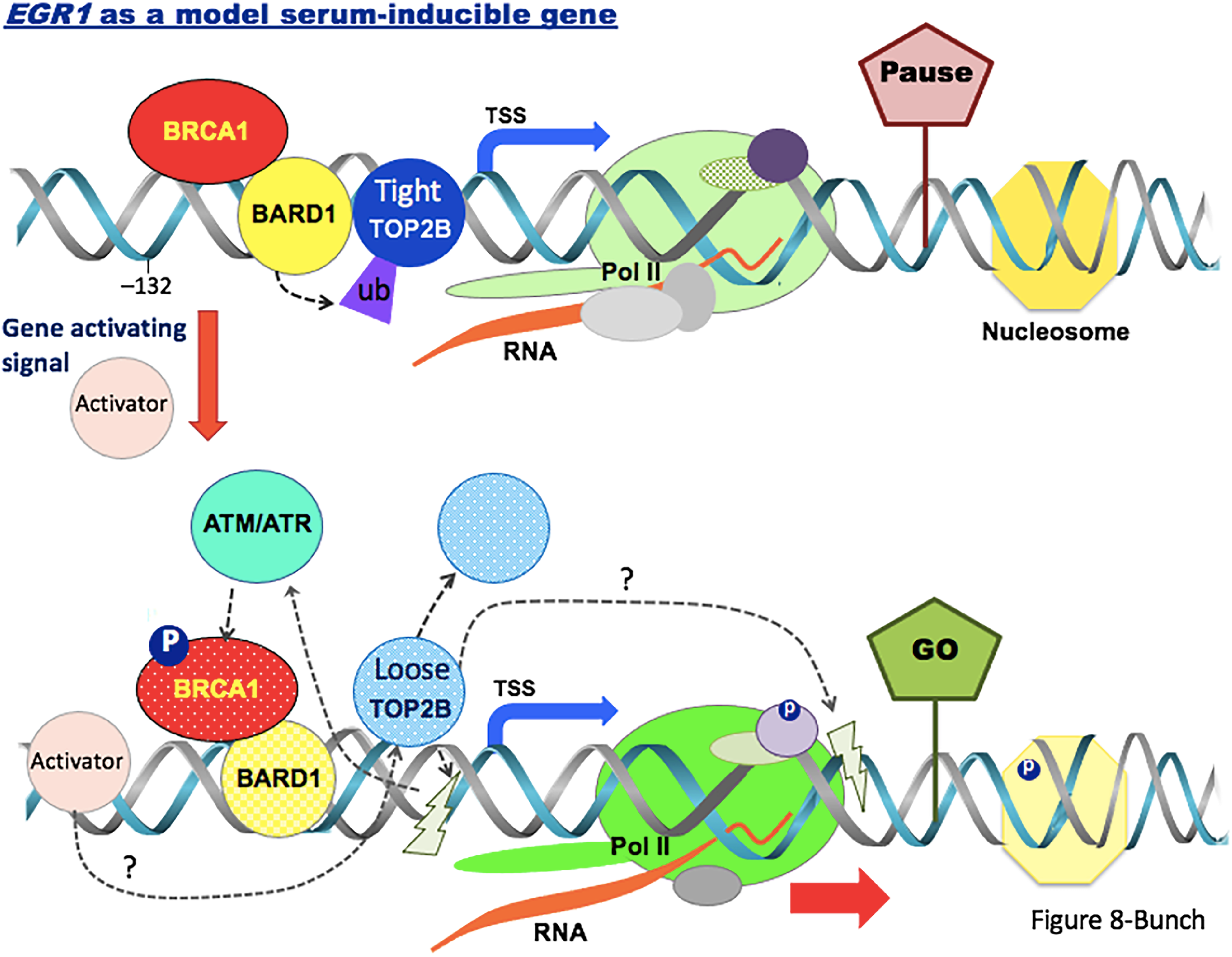
Model of the regulation of BRCA1-BARD1 complex-mediated TOP2B ubiquitination and stability. During the resting state of transcription, Pol II is paused in the promoter-proximal site in hIEGs. The pausing is induced and stabilized by various factors including transcription factors, nucleosome modifiers, and nucleic acids. BRCA1-BARD1 complex is engaged with hIEGs and binds to and ubiquitinates TOP2B in the *EGR1* TSS. This ubiquitination (marked as ub) confers an enhanced DNA binding affinity on TOP2B (Tight TOP2B). In transcriptional activation, ATM/ATR phosphorylates its substrates including BRCA1 at S1524 (phosphorylation, marked as P in a blue circle) and phosphorylated BRCA1 alters the interaction between BARD1 and TOP2B, which mitigates TOP2B ubiquitination. This event destabilizes and dissociates TOP2B (Loose TOP2B) from the TSS. Thus, BRCA1 phosphorylation and the interplay between BRCA1-BARD1 complex and TOP2B appear to be crucial in transcription of stimulus-inducible genes in humans. Lightening marks are proposed catalysis sites of TOP2B based on our previous and current study (Bunch et al., 2015).

## Discussion

Our findings suggest that the BRCA1-BARD1 complex is important for serum-inducible transcription. We showed that BRCA1 located in the TSS is phosphorylated at S1524 during transcriptional activation for representative serum-inducible genes *JUN, FOS, MYC*, and *EGR1* (**Fig. 1, 2**). BRCA1 binding to the DNA might be a weak or indirect one, perhaps mediated through other proteins such as BARD1 because it was barely detectable in the immobilized template assay. The phosphorylation of BRCA1 S1524, mediated by ATM and ATR, is important for the active gene expression of *EGR1 in vivo* and *in vitro* (**Fig. 1–3**). Mechanistically, we suggest that BARD1 physically interacts with and ubiquitinates TOP2B (**Fig. 5a**). The ubiquitination state of TOP2B is a key to modulate its stability and DNA association (**Fig. 5b, 6d, 7a–f**), and the BRCA1-BARD1 complex functions as an E3 ligase for TOP2B (**Fig. 5b–e**). The functional interaction between the BARD1 and TOP2B appears to be regulated by BRCA1 phosphorylation at S1524 (**Fig. 5a**). Furthermore, the phosphorylation of BRCA1 S1524 controls the association between TOP2B and DNA by modulating the interaction between BARD1 and TOP2B (**Fig. 5a, 7e**). Unphosphorylatable BRCA1 mutated at S1524 augments TOP2B ubiquitination and strengthened TOP2B binding to the *EGR1* TSS (**Fig. 5a–e, 7e**), suggesting a tighter DNA binding by ubiquitinated TOP2B during the resting state of transcription or RNA Pol II pausing.

In addition, we identified the E2 enzymes, UBCH3, 6, and 5a–c, required for the BRCA1-BARD1 complex to ubiquitinate TOP2B through the *in vitro* biochemical assays (**Fig. 5e**; **Supplementary Fig. 5F, 6A**,**B**). In particular, UBCH3 and UBCH6 together with the BRCA1-BARD1 yielded discretely shifted bands of ubiquitinated TOP2B (**Fig. 5e**). UBCH3 is known to regulate cell cycle for the entry into the S phase (King et al., 1996) and UBCH6 monoubiquitinates H2B during gene activation (Zhu et al., 2005). On the other hand, UBCH5 enzymes, known to poly-ubiquitinate proteins including p53 and BRCA1 (Brzovic et al., 2006; Scheffner et al., 1994), appear to poly-ubiquitinate TOP2B (**Fig. 5e**). Further study is required to determine the roles of these E2 enzymes for the BRCA1-BARD1 complex-mediated TOP2B ubiquitination.

Genomic analyses showed that BRCA1 colocalizes with TOP2B in a large number of genes including serum-inducible genes (**Fig. 3f, 4**). Due to the resolution of conventional ChIP-seq, the exact TOP2B binding and catalysis sites in the TSS remain of great interest. We could map the binding site of TOP2B with a higher affinity (–132 to +62) in the *EGR1* TSS and showed that BARD1 is colocalized with TOP2B within the region (–132 to –15, **Fig. 6a–d, 7a–c**). It is still unclear whether the TOP2B-DNA resolution site(s) is/are limited to the TSS or is/are extended to the gene body, along with transcriptional activation and elongation, questions which await further examination. In fact, TOP2B and TOP2B-mediated DDR signaling during transcription occurs in the entire promoter, TSS, and gene body of a transcriptionally activated gene (Bunch et al., 2015). In addition, a recent genomics study found that Top-DSB is mapped in the promoter along with CTCF and in the promoter-proximal/gene body (Gittens et al., 2019) whereas another study without etoposide poisoning mapped TOP2B binding sites to the promoter (Canela et al., 2017; Madabhushi et al., 2015; Uuskula-Reimand et al., 2016). Interestingly, the Top-DSB sites in the TSS and gene body appear to be closely correlated with transcriptional activity and gene expression (Bunch et al., 2015; Gittens et al., 2019). From these earlier and more recent data, it can be speculated that TOP2B is mainly associated with the promoter of a gene during the resting stage of gene expression, while its function might become extended to the TSS and gene body along with the promoter region in transcriptional activation. Validation of this hypothesis awaits further study. Because the DNA torsional stress during transcription requires TOP2B-mediated DSB for resolution, the fidelity of the enzyme for removing DNA supercoiling and immediately resealing the broken DNA ends is thought to be crucial. A recent study suggested that TOP2A, not TOP2B, is responsible for regulating supercoiling without provoking DSB in the promoter of hIEGs and inhibiting TOP2A activates transcription (https://www.biorxiv.org/content/10.1101/2020.05.12.091058v1.full). This may suggest a potential competition or redundant function of TOP2A and TOP2B in the TSS while TOP2B functions both TSS and gene body in hIEGs. Other studies have indicated the frequent and spontaneous formation of TOP2B-DNA adducts through abortive catalysis during the transcription of stimulus-inducible genes under normal physiological conditions (Morimoto et al., 2019; Sasanuma et al., 2018), proposing the prompt release and precise repair of TOP2B-DNA adducts is necessary to preserve genomic stability. Our current work suggests that BRCA1-BARD1 involvement controls TOP2B stability on DNA in transcription. These findings raise important questions regarding not only the regulatory mechanisms of TOP2B activity by BRCA1-BARD1 complex for transcriptional initiation and elongation but also the mechanisms of DNA repair of Top-DSB by this complex.

We propose that TOP2B and BRCA1-BARD1 interaction regulates TOP2B association with or dissociation from the TSSs of stress-inducible genes for the regulation of gene expression (**Fig. 8**). In this model, transcriptional activation and Pol II pause release trigger the phosphorylation of BRCA1 by the PI3 kinase family proteins, ATM and ATR. Phosphorylated BRCA1 weakens the functional interaction between BARD1 and TOP2B, which reduces TOP2B ubiquitination and in turn loosens TOP2B from the promoter of the activated gene. In fact, there are more residues in BRCA1 that are reportedly phosphorylated by ATM and ATR upon DNA damage, including Ser1423 (Gatei et al., 2001). In this context, BRCA1 SA or SD substitutions in the current study may have exhibited partial effects of phosphorylation of BRCA1, and further *in vitro* and *in vivo* studies may be required to examine the full function of BRCA1 phosphorylation in TOP2B regulation. We suggest that BRCA1-BARD1-mediated ubiquitination of TOP2B is modulated by BRCA1 in a phosphorylation dependent manner, and ubiquitinated TOP2B binds to the DNA with greater stability than without it. The phosphorylation of BRCA1 at S1524 by ATM and ATR appears important for discouraging TOP2B ubiquitination, thus facilitating the release of TOP2B from the DNA. Our biochemical analyses demonstrated that TOP2B is bound to the *EGR1* promoter and TSS, along with BARD1. Combining with our previous study in which DNA strand break and activated DNA-dependent protein kinase catalytic subunit (DNA-PKcs, phosphorylated at T2609) were observed in the TSS near the Pol II pausing sites of stimulus-inducible genes upon transcriptional activation (Bunch et al., 2015), less ubiquitinate, loosened TOP2B might mediate DNA break in the promoter and downstream of TSS (**Fig. 8**). Our findings indicate a novel and important physiological role for BRCA1-BARD1 complex and the mechanism by which the complex modulates TOP2B in transcription. In addition, it should be noted that DSBs are, in general, repaired more effectively in transcriptionally active units (Aymard et al., 2014; Puc et al., 2017). For the novel function of the BRCA1-BARD1 complex in Pol II transcription, we propose that BRCA1 activation during gene expression may be involved in the regulation of DNA break and repair to ensure genome integrity, which awaits future investigations.

## Acknowledgments

We thank D.J. Taatjes at the University of Colorado for providing HeLa nuclei and technical and material support, and we thank M. Erdos and L. Brody at the United States National Institute of Health (NIH) for providing pcBRCA1-385 and information related to the construct. We appreciate D. Levine at NIH for the helpful discussion and perspectives and for the interest expressed in our research. We thank T. Volkert and S. Gupta in the Whitehead Institute Genomic Technology Core and B.P. Lawney and Y.E. Wang at the Quantitative Biomedical Research Center of the Harvard T.H. Chan School of Public Health for providing with the generation and sequencing of ChIP-seq libraries, bioinformatics, and technical support. We are grateful to S. Lee, J. Heo, S. Jeon, J. Sohn, H. Choe, K. Baek, S. Heo, and all members of Bunch Lab at Kyungpook National University for their technical assistance. H.B. thanks S. Buratowski at Harvard Medical School and D.J. Taatjes at the University of Colorado for research advice and J. Christ, G.P. Hugenberger, J. Park, H. Yang, and John and D.Y. Bunch for their loving encouragement throughout the course of this work. This research was supported by grants from the National Research Foundation (NRF) of the Republic of Korea (2017R1D1A1B03030548) to H.B., from the National R&D Program of the Cancer Control, Ministry of Health & Welfare of the Republic of Korea (1720100) to K.K., from the NRF, Ministry of Science, and ICT of the Republic of Korea (2017R1A2B4005501) to D.-H.C., Mayo Clinic startup funds to M.J.S., and from the NIH of the USA (CA233594) to B.P.C.

## Author Contributions

JJ carried out molecular cloning, protein purification, template DNA preparation, immunoblotting, *in vitro* ubiquitination, and EMSA. HB performed the protein purification, immunoprecipitation, immunoblotting, *in vitro* ubiquitination, and immobilized template, *in vitro* transcription, ChIP, and quantitative real time PCR assays. KK and BPCC performed bioinformatics. ATC and MJS prepared proteins from human cells and performed prerequisite functional analysis with them. DJ and DHC carried out immunofluorescence experiments. DK performed cloning, quantitative PCR and protein purification. SKC advised in the interpretation of the data. HB designed the experiments, analyzed the data, and wrote the manuscript.

## Data and materials availability

All data are available in the manuscript or the supplementary material.

